# Using structural connectivity to augment community structure in EEG functional connectivity

**DOI:** 10.1101/831743

**Authors:** Katharina Glomb, Emeline Mullier, Margherita Carboni, Maria Rubega, Giannarita Iannotti, Sebastien Tourbier, Martin Seeber, Serge Vulliemoz, Patric Hagmann

## Abstract

Recently, EEG recording techniques and source analysis have improved, making it feasible to tap into fast network dynamics. Yet, analyzing whole-cortex EEG signals in source space is not standard, partly because EEG suffers from volume conduction: Functional connectivity (FC) reflecting genuine functional relationships is impossible to disentangle from spurious FC introduced by volume conduction. Here, we investigate the relationship between white matter structural connectivity (SC) and large scale network structure encoded in EEG-FC. We start by confirming that FC (power envelope correlations) is predicted by SC beyond the impact of Euclidean distance, in line with the assumption that SC mediates genuine FC. We then use information from white matter structural connectivity (SC) in order to smooth the EEG signal in the space spanned by graphs derived from SC. Thereby, FC between nearby, structurally connected brain regions increases while FC between non-connected regions remains unchanged, resulting in an increase in genuine, SC-mediated FC. We analyze the induced changes in FC, assessing the resemblance between EEG- and volume-conduction-free fMRI-FC, and find that smoothing increases resemblance in terms of overall correlation and community structure. This result suggests that our method boosts genuine FC, an outcome that is of interest for many EEG network neuroscience questions.

**Author summary:** In this study, we combine high-density EEG recorded during resting state with white matter connectivity obtained from diffusion MRI and fiber tracking. We leverage the additional information contained in the structural connectome towards augmenting the source level EEG functional connectivity. In particular, it is known - and confirmed in this study - that the activity of brain regions that possess a direct anatomical connection is, on average, more strongly correlated than that of regions that have no such direct link. We use the structural connectome to define a graph and smooth the source reconstructed EEG signal in the space spanned by this graph. We compare the resulting “filtered” signal correlation matrices to those obtained from fMRI and find that such “graph filtering” improves the agreement between EEG and fMRI functional connectivity structure. This suggests that structural connectivity can be used to attenuate some of the limitations imposed by volume conduction.

## Introduction

Electroencephalography (EEG) measures neural signals directly (Buzsáki et al., 2012) on a time scale of milliseconds, fast enough to be relevant for neural events. In MEG and fMRI, the study of functional connectivity (FC) with the tools of whole-brain network neuroscience has yielded a multitude of important and interesting insights (see Bassett & Sporns, 201), for a review). Concurring findings show that FC between regions of interest (ROIs)/sources located in the gray matter is in part shaped by anatomical connections of the structural connectivity (SC; obtained from dMRI and fiber tracking), such that the strength of SC (fiber count, density) is predictive to some degree of the strength of FC (correlation, coherence, etc.; Abdelnour et al., 2018; Atasoy et al., 2016; Cabral et al., 2014; Damoiseaux & Greicius, 2009; Deco et al., 2013; Glomb et al., 2017; Goñi et al., 2014; Hagmann et al., 2008; Honey et al., 2009; Meier et al., 2016; Tewarie et al., 2014, 2019; Vincent et al., 2007). This finding has been shown to extend to EEG data on the source level (i.e., signals recorded on the scalp projected into the gray matter) using analytical (Chu et al., 2015; Wirsich et al., 2017) and modelling approaches (Bhattacharya et al., 2011; de Haan et al., 2012; Finger et al., 2016; Pons et al., 2010; Ponten et al., 2010; van Dellen et al., 2013). The main hurdle when trying to understand EEG network architecture and -dynamics is signal leakage due to volume conduction, which obscures genuine functional relationships between sources in the brain: Additionally to the low spatial resolution (Buzsáki et al., 2012; Schoffelen & Gross, 2009; Srinivasan et al., 2007) and SNR intrinsic to the EEG signal, the interaction of the electric field with the tissue creates “sham” functional connections whose strengths depend on the Euclidean distance between locations. In order to circumvent these problems, it has been suggested that zero-lag statistical dependencies should be removed altogether from FC analysis since signal leakage is instantaneous, resulting in measures such as imaginary coherence (Nolte et al., 2004) and phase lag index (Stam et al., 2007). Contrary to this, it has been pointed out that zero-lag statistical dependencies still carry meaningful information about ongoing activity (Pascual-Marqui et al., 2017; Tognoli & Kelso, 2009; Uhlhaas et al., 2009), and may therefore be particularly important for data recorded during resting state. Furthermore, approaches exist that orthogonalize the time series, removing common dependencies between sources (Brookes et al., 2012; Colclough et al., 2015; Hipp et al., 2012; Wens et al., 2015).

In this study, we propose an approach that incorporates additional information from SC into EEG-functional connectivity on the source level.

We first replicate the previous finding (Chu et al., 2015; Finger et al., 2016; Siems et al., 2016; Wirsich et al., 2017) that SC partially shapes FC in EEG, beyond the impact of Euclidean distance induced by volume conduction. Subsequently, we use additional information in the SC to augment the EEG functional signal by applying a low-pass filter (or smoothing procedure) in the space spanned by the SC-graph. The underlying motivation is that both SC and FC decay with increasing Euclidean distance. From an evolutionary standpoint, it makes sense for functionally related regions to be close together (Tomasi et al., 2013). As a result, it is impossible to disentangle the contributions of volume conduction and genuine FC mediated by SC to a statistical dependency measured between two brain regions. However, if one assumes that genuine FC is mediated by SC, and therefore, genuine FC is higher between brain regions that are directly anatomically connected, increasing the impact of SC would also increase the contribution of genuine FC relative to FC generated by volume conduction. A similar approach has previously been shown to improve source estimation in EEG (Hammond et al., 2013). We use an fMRI-FC acquired completely independently of the EEG dataset in order to determine whether our filtering procedure leads to improvements in terms of macroscopic network structure encoded in the FC. Numerous studies have shown that the macroscopic network structure is, on a coarse level, similar between fMRI and EEG (Britz et al., 2010; Coito et al., 2019; Liu et al., 2018; Musso et al., 2010). These networks have been demonstrated to be relevant on the several spatial and temporal scales of different recording techniques (Brookes et al., 2011; Kucyi et al., 2020; Liu et al., 2017). We hypothesize that, if our filtering procedure indeed strengthens genuine FC mediated by SC, we should see that the network structure encoded in the EEG-FCs becomes more similar to that encountered in fMRI, a recording technique that is not impacted by volume conduction; we quantify this by testing whether the EEG-FCs computed from filtered time courses are more similar to fMRI-FC than the original, unfiltered signals, both overall and by explicitly analyzing the FC-matrices’ community structure. Our results suggest that incorporating information from the SC by means of graph filtering leads to a large-scale network structure in EEG-FC that is more similar to known canonical resting state networks, making our approach a possible alternative to other methods that aim at correcting for volume conduction, especially for the study of large-scale functional networks.

## Results

### SC provides additional predictive power for EEG-FC

Our goal is to use the structural connectivity (SC) matrix to boost functional connectivity (FC) that is mediated by white matter anatomical connections in source-level EEG resting state time series recorded from N=18 subjects (see Figure 1 for an illustration of our approach).

**Figure 1:**
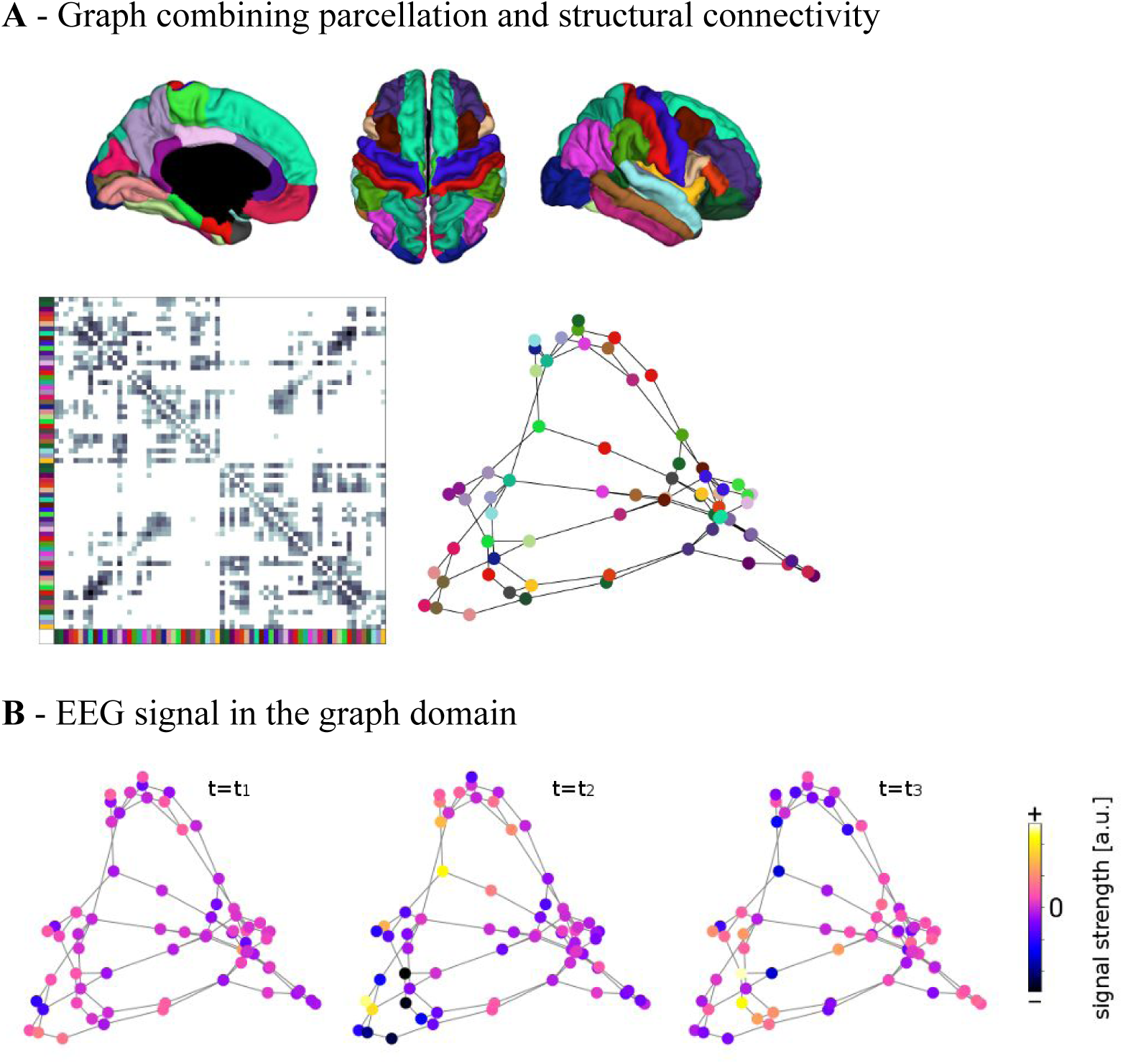
**A** - Brain regions are given by the Lausanne 2008 parcellation (top row). Diffusion MRI and fiber tracking reveal fiber bundles that exist between these regions, and are summarized in the structural connectivity matrix (bottom left; brain regions from above are color coded). The graph used for filtering is defined by nodes corresponding to the parcellation’s brain regions (color coded) and edges corresponding to the fiber bundles in the SC matrix (bottom right; lengths of edges are approximately inversely proportional to the weight in the structural connectivity matrix). **B** - The graph defined by parcellation and SC does not change over time. EEG signals that do vary over time are conceptualized as activation strengths of the nodes of the graph. Signals are thought to propagate along the fiber bundles, i.e. the edges, to neighboring nodes.

One of our main assumptions is that genuine FC is in part mediated by SC (Chu et al., 2015; Finger et al., 2016; Wirsich et al., 2017). To test this, we first show that SC can predict FC beyond the common dependence of FC and SC on Euclidean distance. We test this assumption by fitting a stepwise general linear model (GLM) in order to quantify how well the following measures predict EEG-FCs computed as envelope correlations from three typical EEG-frequency bands (alpha: 8-13 Hz, beta: 13-30 Hz, gamma: 30-40 Hz), averaged over all subjects:

1. SC in the form of search information (Goñi et al., 2014), referred to as SC_SI_, a measure that is derived from fiber counts and that is non-zero for all connections, yielding a connectivity matrix that is dense just like the FC matrix (Figure S1); the intuitive interpretation of search information is that it measures how “hidden” the shortest path between two ROIs is. Note that search information and fiber count are roughly *inversely* proportional.
2. Euclidean distance (ED) between ROI centers,
3. relative regional variance (i.e. the variance of each ROI time course, normalized such that the maximum variance in each subject equals 1) as an estimate of SNR,
4. ROI size (number of voxels in the parcellation).

The 3rd and 4th predictors are control variables for possible confounds. Note that in each case, we predict a dependent univariate variable - the EEG-FC - with an independent univariate variable, such that each pair of brain regions is a sample (i.e. we have (*N* * *N* − *N*)/2 samples). We analyze how well the independent variables can predict the FC in two different ways: On the one hand, we use each variable as the only predictor variable (plus intercept; blue bars in Figure 2; “single variable model”). On the other, we test their predictive power when they are entered progressively into a GLM that includes first order interaction terms (orange curves in Figure 2; “full model”). The latter case allows us to quantify the additional predictive power that each variable has, given all other predictors. This is important because the predictors are not independent of each other; most prominently, there is a three-way-dependence between SC, Euclidean distance, and FC, such that both SC and FC decay with increasing Euclidean distance. Figure 2A shows the results in terms of explained variance for the EEG-FCs and for comparison, for an fMRI-FC (average pairwise correlations over 88 subjects, see Methods for details; see SI Table 1 for detailed results of the GLM analysis). The correlation between the average EEG-FCs and the fMRI-FC is around 0.50 for all three bands (alpha, beta, gamma).

**Figure 2:**
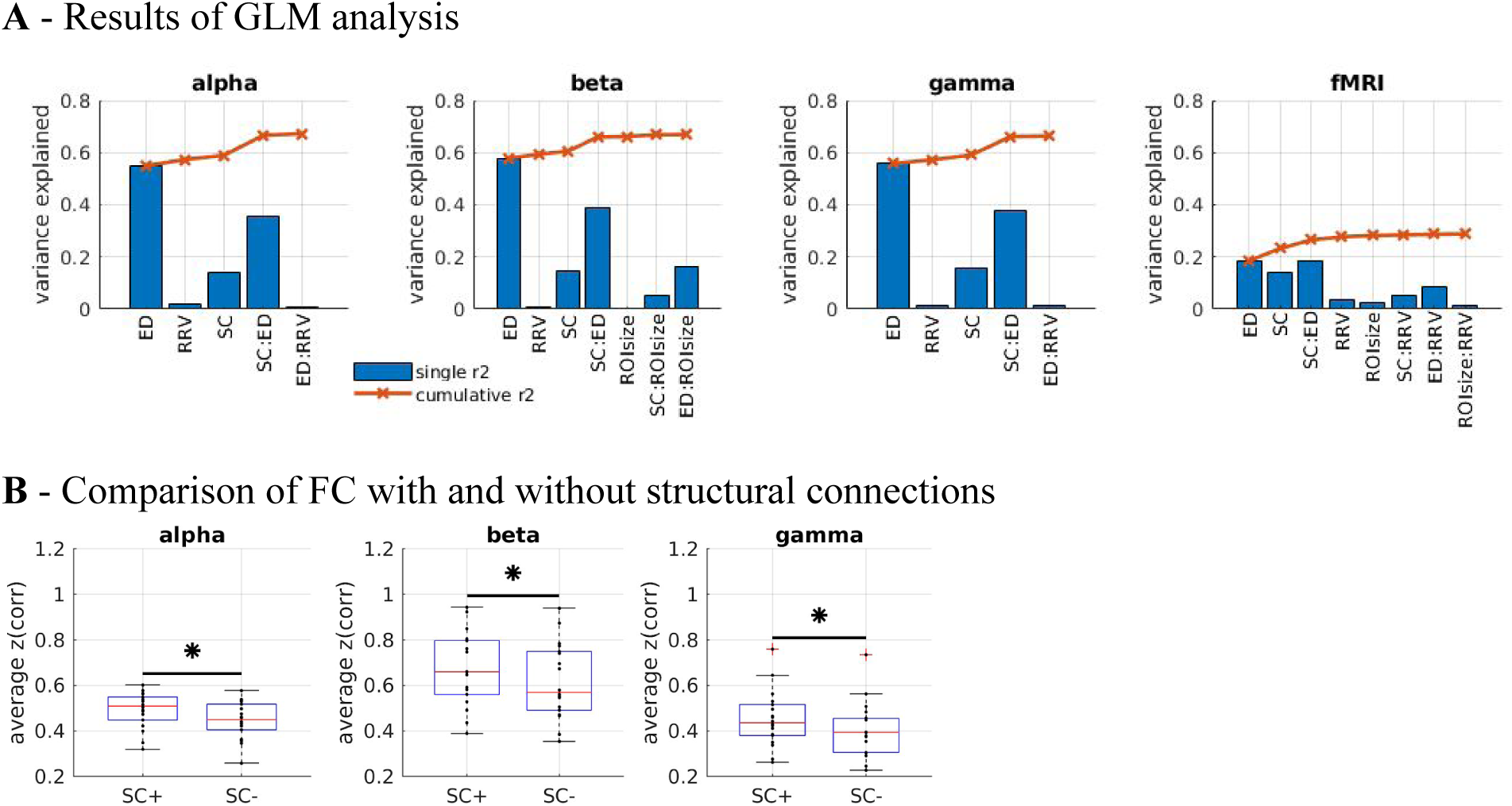
**A** - Stepwise general linear model results for EEG alpha, beta and gamma band, and for fMRI, using Euclidean distance (ED), relative regional variance (RRV), structural connectivity (SC; search information), and ROI size (number of voxels) as predictors. Only significant predictors are shown, in the order in which they were entered into the model (see Methods for details). Blue bars: variance explained when variables are used as the only predictor in separate “single variable” GLMs. Orange curves and crosses: Cumulative explained variance achieved when using all variables up to the variable corresponding to this data point (i.e. the variable in question and all variables to the left of the data point). **B** - Comparison between average EEG-FC values for pairs that are connected by SC (“SC+”) and those that are not (“SC-”). The samples that are compared are matched in their ED distribution to control for the fact that pairs that are connected tend to be closer together than those that are not. Stars mark significant differences according to the Wilcoxon signed-rank test at alpha=0.05 (Bonferroni-corrected for multiple comparisons).

Figure 2A shows that ED is the strongest predictor in both EEG and fMRI. This is true both for the single variable models (greatest explained variance as indicated by blue bars) and the full model (as indicated by the fact that they are the variables that are entered first into the model). However, the variance explained by ED is much higher in EEG than in fMRI, namely 0.55 in EEG (alpha band; other bands similar, full list in Table S1) and 0.18 in fMRI. This is due to the effect of volume conduction which introduces spurious correlations dependent on ED in the case of EEG-FC.

When using SC_SI_ as the only predictor, we find that the dependency of FC on SC_SI_ is very similar in both modalities (r^2^ of SC_SI_ alone 0.14 in both modalities). In both cases, FC values are high between close-by pairs of brain regions (small ED; see Figure S2A). For fMRI, there are also highly correlated pairs that are separated by an intermediate distance, and close-by pairs that are barely or not at all correlated. In contrast, for EEG, all far-away pairs of ROIs have low correlations and all close-by pairs have high correlations. We checked for which connections the prediction of FC by SC_SI_ was worst, i.e. had the largest residuals (Figure S2B). We found that for fMRI, the largest errors occur on the secondary diagonal, replicating the well-known result that interhemispheric connections are underestimated in the SC. In EEG, this does not contribute as much to the unexplained variance as the FC between homotopic regions is low compared to FC between close-by pairs of regions. In summary, while the variance explained by the SC_SI_ is the same in both modalities, the structure of this dependency is different. In both cases, SC_SI_ explains an additional 3-4% of the variance (SI Table 1) after regressing out Euclidean distance. For fMRI, some connections with intermediate distances remain underestimated after adding SC_SI_ as a predictor, again related to interhemispheric connections (Figure S2B, right panel). For EEG, the most severely underestimated FC values are related to small distances, indicating that neither Euclidean distance nor SC_SI_ can by themselves account for some of the large EEG-FC values between nearby pairs of regions. Note that the actual contribution of SC_SI_ is likely to be higher than 3-4%, as SC strength is itself dependent on Euclidean distance (two regions that are close together are more likely to be connected by white matter tracts; furthermore, short tracts are more easily traced by fiber tracking algorithms, Jones, 2010). Indeed, due to this mutual dependence, the actual contribution of SC_SI_ cannot be estimated using this approach.

Furthermore, a significant positive interaction term between ED and SC_SI_ contributes to the prediction in both fMRI and EEG. The correlation between these two variables (ED and SC_SI_) is 0.48, in line with previous findings (Wirsich et al., 2017). Since SC and SC_SI_ are negatively correlated (see Methods), this translates to two interpretations: ED has less of an impact on the FC between ROI pairs that have a strong SC-connection (high weight in the SC matrix); and the strength of the SC connection has less of an impact on the FC between ROI pairs that are far apart from each other (high ED).

A simple prediction from the hypothesis that FC is shaped by SC is that ROI pairs that are connected via white matter tracts should exhibit stronger FC than those that are not (Chu et al., 2015). In order to control for the common dependence of FC and SC on Euclidean distance, we compare average FC values over subsamples of pairs of ROIs that are matched in their Euclidean distance distribution. Figure 2B shows that even in those matched subsamples, there is indeed a significant difference between the mean FC values (Wilcoxon signed-rank test at alpha=0.05, Bonferroni-corrected for multiple comparisons) between structurally connected and unconnected ROI pairs.

### Graph filtering increases resemblance between EEG-FC and fMRI-FC

Building on the finding that SC shapes EEG-FC beyond the impact of ED, we use the fact that the EEG data live on a graph defined by the SC (Figure 1). In the following, we perform spatial smoothing, or low-pass filtering, in graph space. This means that the activity in one node of the graph - an ROI - is smoothed by computing a weighted average of the ROI’s activity and the activity of its nearest neighbors, i.e. nodes with which it is anatomically connected:

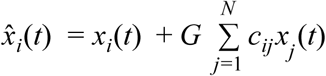

Here, *c*_*ij*_ is the entry in the SC which corresponds to the pair of regions *i* and *j*, and *G* is the scalar “filter weight” which scales how much impact node *i* ’s neighbors *j* have on the activity of *i*. This will increase the effect shown in Figure 2B: Connected nodes’ activity will become more similar and unconnected nodes’ activity will become less similar. Since spurious correlations depend only on Euclidean distance, but not on the SC per se, this would correspond to an increase of genuine FC relative to spurious FC.

We filter the single epoch, source-level activity *x*_*i*_(*t*) of each ROI using the above formula and compute FC matrices from the filtered data 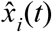 as before (envelope correlations in alpha, beta, and gamma frequency bands). We use four different graphs (Figure 3, from left to right):

**Figure 3:**
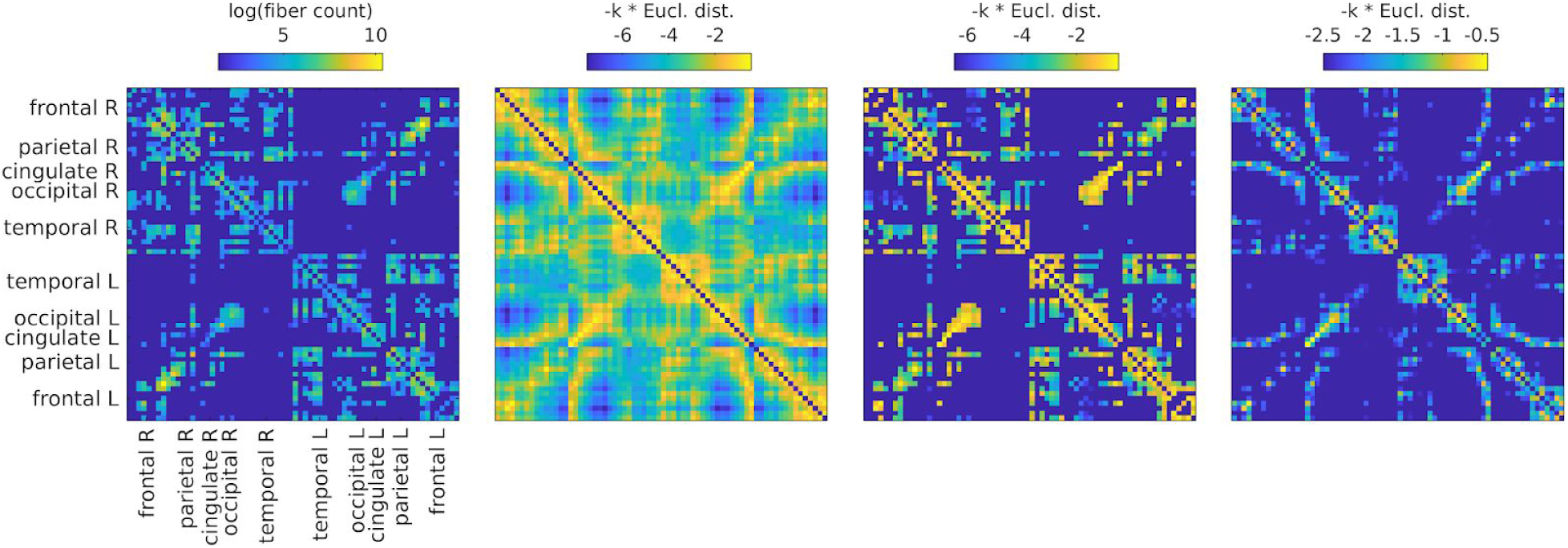
All SCs from which graphs for filtering are derived. From left to right. 1) SC derived from fiber tracking, averaged over subjects (log is used only for visualization purposes); 2) SC derived from Euclidean distances by using the distances as negative exponents; 3) SC derived from Euclidean distances (as 2), but masking the distances by using present connections as in 1); 4) SC derived from Euclidean distances (as 2), but keeping only shortest distances such that the density is the same as in the left and middle right panels.

1. The SC (number of fibers) itself, averaged across subjects according to Betzel et al. (2019). This graph has a connection density of 25%.
2. A graph derived from Euclidean distances, setting connection weights with *exp*(− *k* * *ED*), where the weight *k* just serves to scale the distribution such that the multiplication of the graph weights with the filter weights will result in effective weights in the same range as for the other SCs. This is a dense graph and will be referred to as “ED full”.
3. A graph derived from Euclidean distances, but exactly matching the SC in terms of existing and absent fibers. This means that the connections are the same as for SC 1, but the weights are set as in SC 2 instead of the fiber count. This graph will be referred to as “ED match”.
4. A graph derived from Euclidean distances, but preserving the density of SCs 1 and 3. This means keeping the connections which correspond to the smallest EDs up to a threshold which leads to the same connection density as in SC1 (and SC 3). We include this graph because otherwise, if matrices 1 or 3 outperform matrix 2, this could just be due to the difference in connection density. This graph will be referred to as “ED dens”.

After filtering, EEG-FCs are computed as for the unfiltered data. As expected, the filtered FCs become more similar to their respective SC_SI_ (Pearson correlation between EEG-FCs and SC_SI_), and the difference in average correlation between structurally connected versus unconnected ROI pairs increases (see Figures S3, S4).

To validate our results, we compare our EEG-FCs to fMRI-FC (obtained by averaging over 88 subjects, see Methods for details) by computing the Pearson correlation between the two matrices for each subject. The goal is to assess whether the changes in FC induced by graph filtering result in a change in the network/community structure encoded in the FC which is in line with known functional networks (Britz et al., 2010; Coito et al., 2019; Liu et al., 2018; Musso et al., 2010). We obtain a comparison for each filter weight *G* and each of the four graphs described above and shown in Figure 3.

Figure 4A shows that correlations between the two matrices obtained from the two modalities increase as hypothesized (results shown for beta band, alpha and gamma similar [not shown]). The graph which reaches the highest maximum correlations between EEG- and fMRI-FC is the one in which Euclidean distance and SC are combined by masking the weights derived from Euclidean distances with the non-zero connections given by the SC (“ED match”; Wilcoxon signed rank test comparing the correlation coefficients of each subject, p<0.05 Bonferroni corrected). The mean correlation increases from 0.42 to 0.51 at a filter weight of G=100, corresponding to a 23% increase (increase computed based on the Fisher z-transformed values as shown in Figures 4A and B: 0.46 and 0.56, respectively), while the increase when using the SC itself is only from 0.42 to 0.44 (Fisher z-transformed values: increase from 0.46 to 0.48). Figure 5 shows the original EEG-FC (beta band) and the EEG-FC derived from filtered data with G=100.

**Figure 4:**
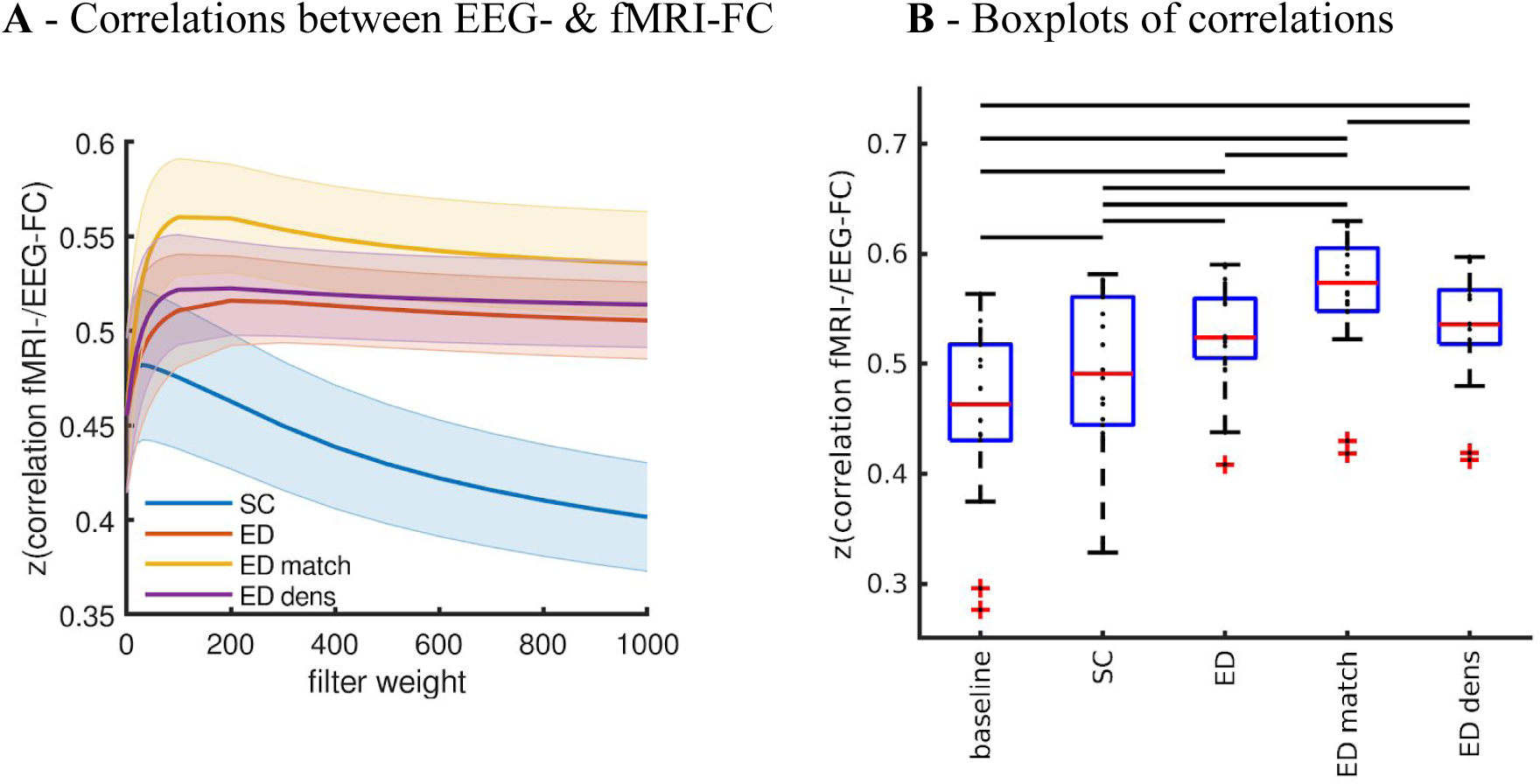
**A**:Fit (z-transformed correlation) between the EEG-FCs (beta band) computed from time courses with different filter weights (G) and the fMRI-FC. The shaded regions mark the 95% confidence interval. **B**: Boxplots summarizing results of the Wilcoxon signed-rank test comparing individuals’ maximum fits (shown in panel A) across versions of the SC as well as to the baseline correlation between unfiltered EEG-FCs and fMRI-FC. Black bars mark significant differences. Red lines mark the median, each black dot marks the value for one subject. Note that we did not compare the medians, but the individual differences (see Methods).

**Figure 5:**
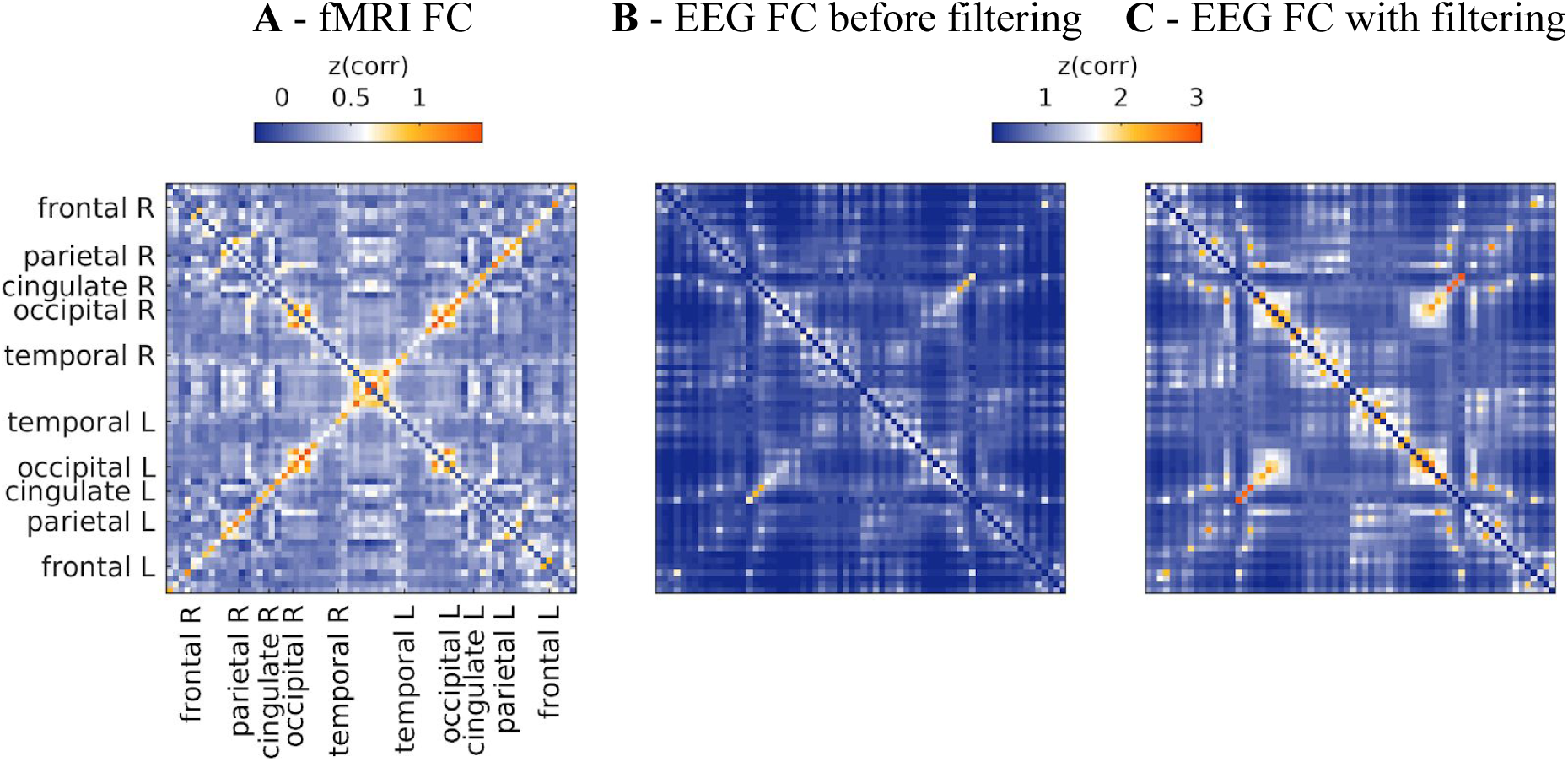
**A**: fMRI-FC. **B** and **C**: EEG-FCs (beta band) before (**B**) and after (**C**) graph filtering with “ED match” and G=100. All correlations are z-transformed.

In order to better interpret these results, we use a control SC in which connections are randomized while preserving the degree distribution and density (Figure S5; see Methods for details). In this case, the fit between EEG- and fMRI-FC increases from 0.42 to 0.44 when using the SC itself, an increase that is almost the same which is achieved with the original matrix, and which remains significant (Figure S5B; Wilcoxon signed rank test, p<0.05 Bonferroni corrected). The remaining three versions of the graph lead to statistically identical increases, i.e. in this case, there is no advantage of using the SC-derived mask on the Euclidean distances (“ED match”) over using the full set of Euclidean distances (“ED full”) or the shortest Euclidean distances only (“ED dens”). Note that, since randomizing the Euclidean distances is not readily possible (as the resulting geometry would need to be consistent), “ED dens” and “ED full” are identical for the randomized and the original SC.

Taken together, increases in fit due to the SC alone can mostly be attributed to the degree distribution (which is preserved in the random SC). Using a graph in which the real SC is used to mask Euclidean distance-derived weights leads to a significant advantage compared to purely Euclidean distance-derived graphs.

We further repeat the analysis using FC matrices computed from white Gaussian noise (WGN-FCs; see Methods) in order to test in how far our results can be explained purely by linear dependencies imposed by the graphs used as filters. We find that the correlation between WGN-FCs that were filtered with the SC and the fMRI-FC reaches a maximum of 0.26 (Figure S6A), indicating that filtering white noise with the SC does not explain the association between fMRI-FC and EEG-FCs, where the correlation is 0.42 without any filtering (Figure 4A). The correlation between WGN-FCs and fMRI-FC reaches a value of r=0.49 (z-transformed value: 0.53) when using only Euclidean distances (dense ED) with G=500. This is comparable to the optimal fit of r=0.51 (z-transformed value: 0.56) obtained with empirical EEG-FCs at G=100 (ED match), but at G=100, the empirical EEG-FCs clearly outperform the WGN-FCs (r=0.35). We also checked whether the fit to fMRI can be explained by the fact that EEG-FCs become more similar to WGN-FCs as the filter weight is increased. At G=500, the filtered EEG-FCs are very similar to the FCs obtained from filtered WGN (average r=0.91 [z-transformed value: 1.5], Figure S4B). At G=100, this correlation is r=0.69 for ED match (z-transformed value: 0.85). As a further comparison, we use the orthogonalization approach described in Colclough et al. (2015) to correct for leakage in the unfiltered data (Figure S7). We found that the correlation between fMRI- and EEG-FCs decreases for each subject (Table S3). Furthermore, we repeat our analyses using coherence and imaginary part of coherence (Figures S8 and S9), the latter of which is assumed to remove zero-lag correlations resulting from volume conduction (Nolte et al., 2004). We found no advantage of these measures over correlations between power envelopes. Specifically, the correlation between fMRI-FC and EEG-FCs computed using this measure was not higher than when using the power envelope correlations (Figure S8A).

### High-quality SCs are necessary to achieve an increase in fit

We perform the same analysis with two additional SC matrices from different cohorts (see Methods for details). The first one is obtained from a small cohort with relatively lower quality than the “primary” SC used in the main analysis above, i.e. DTI was used instead of DSI, as well as a smaller number of subjects (N=20). The second consists of the 45 subjects of the Human Connectome Project’s (HCP) “retest” dataset, data of very high quality. When using the lower quality data (Figure S10), there is no additional benefit of using the SC matrix as a mask on the matrix of Euclidean distances (fit between EEG- and fMRI-FC is the same for “ED full” and “ED match”). However, when using the high quality HCP SC matrices (Figure S11), ED match performs significantly better than the three other graphs, as is the case for the primary SC matrix. This also shows that our results do not depend on the fMRI- and the dMRI-data stemming from the same subjects.

Figure S12A illustrates that the primary matrix used in this study as well as the HCP matrix possesses a higher density of interhemispheric connections than the DTI-derived matrix (primary: 12%, HCP: 20%, DTI: 9%, see Figure S12). This may be due to the fact that interhemispheric fibers are found more consistently in the DSI-based datasets (primary and HCP) than in the DTI-based one, but could also be attributed to variations in the tractography techniques (deterministic vs probabilistic, seeding from white matter versus seeding from gray matter/white matter interface). The absence of some interhemispheric connections leads to high errors in the resulting filtered EEG-FC for the DTI SC-matrix (Figure S12B). This shows that our method relies in part on the improvement of interhemispheric FC, an effect that can only be achieved with high-quality diffusion data.

### Graph filtering increases resemblance between EEG- and fMRI-community structure

In the following, we explore the effect of graph filtering on the EEG-FC structure. To this end, we use FCs averaged over all subjects. First, we consider seed correlations. In order to make correlations comparable, we resample average FCs such that FC values are normally distributed around mean 0 and with a standard deviation of 1. Figure 6 shows the normalized correlations between two seed regions and all other ROIs in the parcellation (i.e. one row/column of the EEG-FC) before and after filtering with the best SC identified above (“ED match”, G=100; beta band). We chose these regions because they exhibit the largest overall change in connection weights with other regions (all changes in Figure S13). In both cases, the correlations to the corresponding region on the other side of the brain are increased.

**Figure 6:**
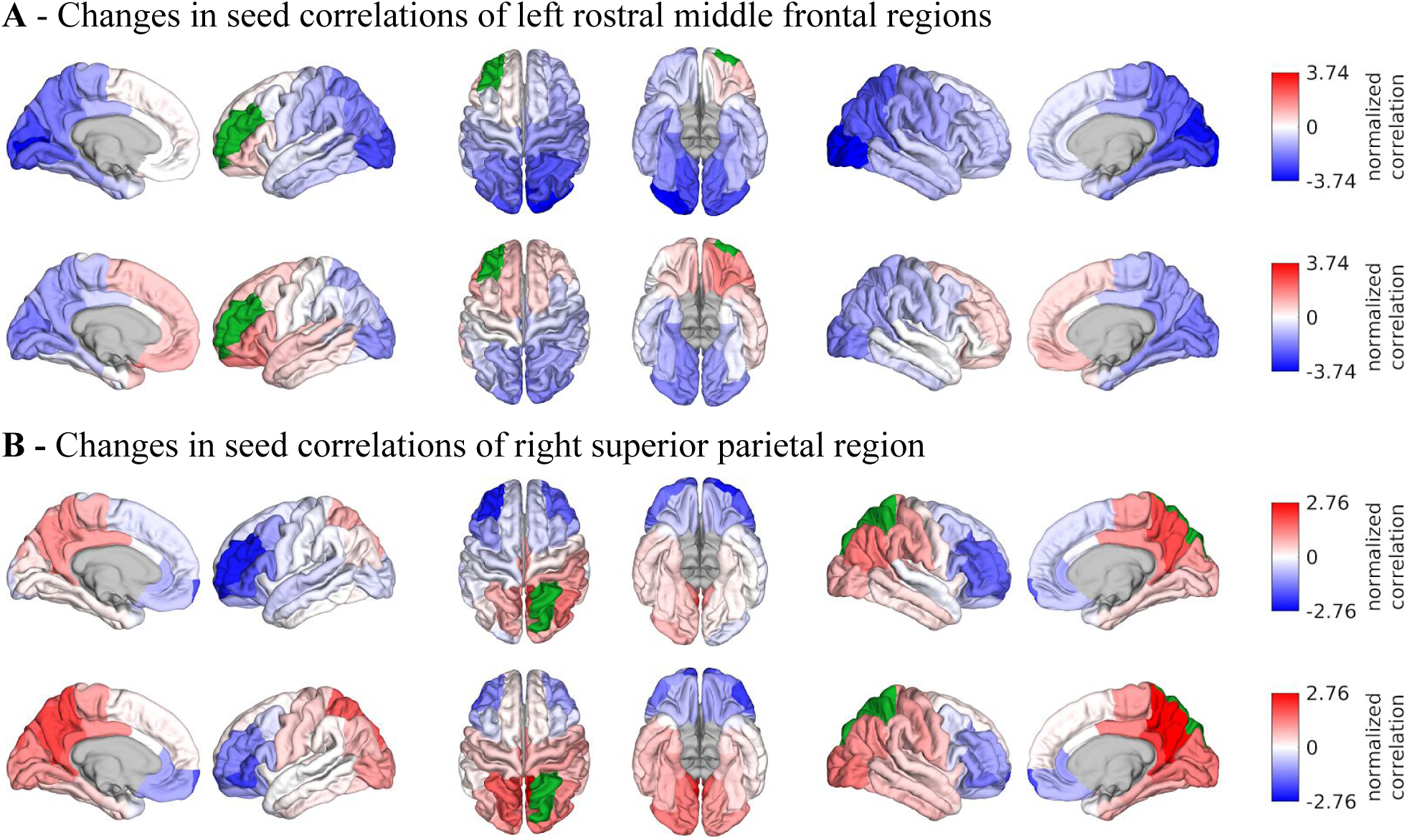
Surface renderings of FC before and after filtering. EEF-FC (beta band; correlation) values were resampled from a standard normal distribution in order to linearize them and make them comparable. **A**: Seed correlations of region left rostral middle frontal before (top panel) and after (bottom panel) graph filtering. **B**: Same as A, but for region right superior parietal.

In general, the correlation structure of many regions changes considerably (Figure S13), but these changes are hard to interpret on a ROI by ROI basis. Therefore, to investigate this further, we extract the community structure of the average EEG-FCs before and after filtering by assessing the probability of any two regions to be assigned to the same community (Figure 7, top row and bottom left). We apply the same procedure to the average fMRI-FC (Figure 7, bottom right). As the filter weight (G) increases, more values in the “community matrices” tend towards 0 or 1, indicating that the variability in clustering outcomes decreases. Interestingly, also the community assignments derived from the fMRI-FC show some variability, especially for certain frontal and temporal regions. Figure 8A shows one example of community assignments. The main difference between EEG and fMRI is that EEG community structure is dominated by the lobe architecture, whereas in fMRI, we see the robust clustering of frontal with middle temporal and posterior cingulate cortices, reminiscent of the default mode network (Laird et al., 2009), as well as the symmetry across hemispheres between temporal regions.

**Figure 7:**
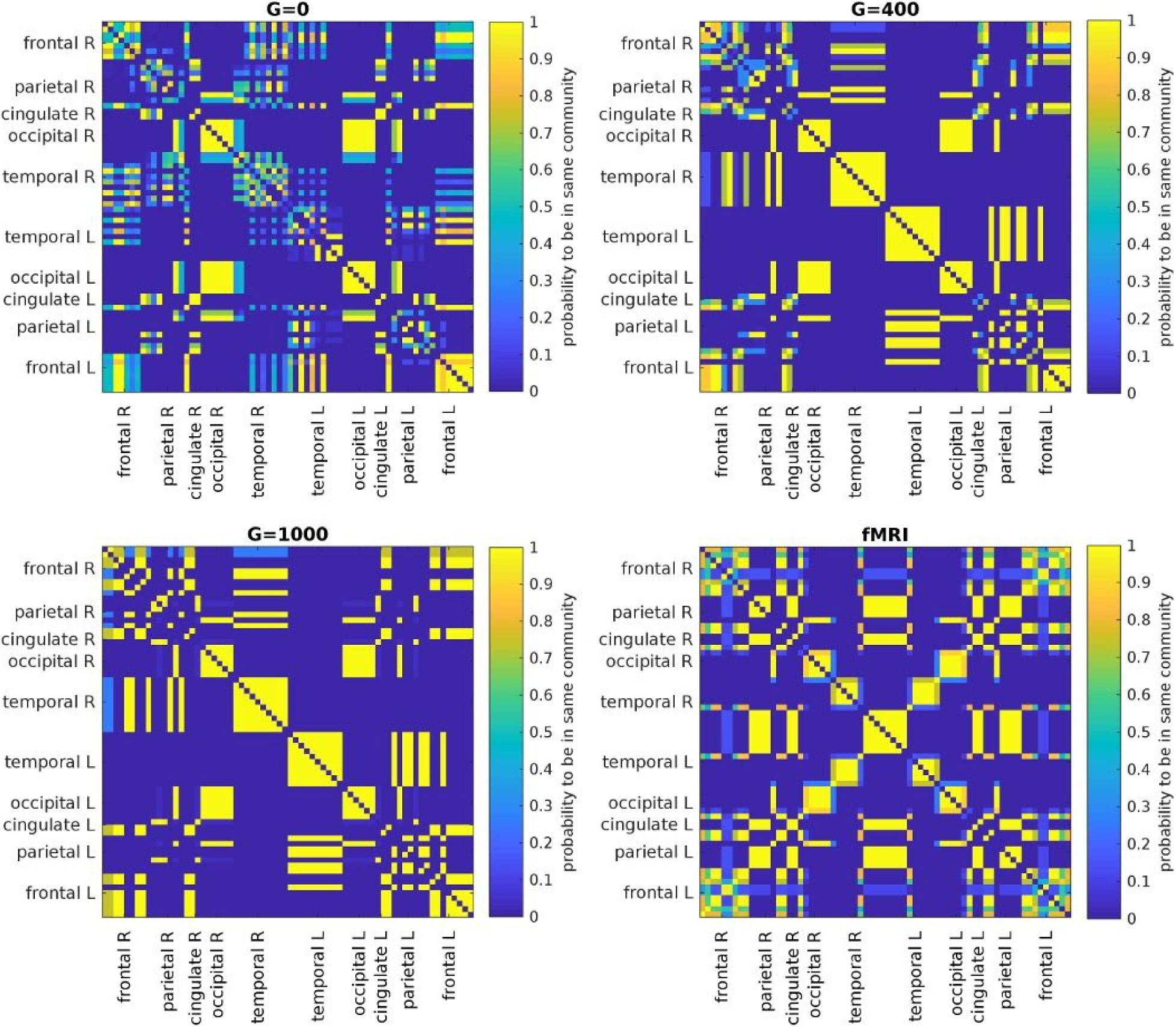
Results of Louvain clustering (“community matrices”). Each matrix shows, for each pair of ROIs, the fraction of repetitions of Louvain clustering (200 rounds, ɣ=1.1) which assigned both ROIs to the same community. ɣ=1.1 and G=400 (upper right matrix) are the parameter setting from a region of the parameter space where the agreement between EEG- and fMRI-community structures were found to be maximal (Figure 9; main text).

**Figure 8:**
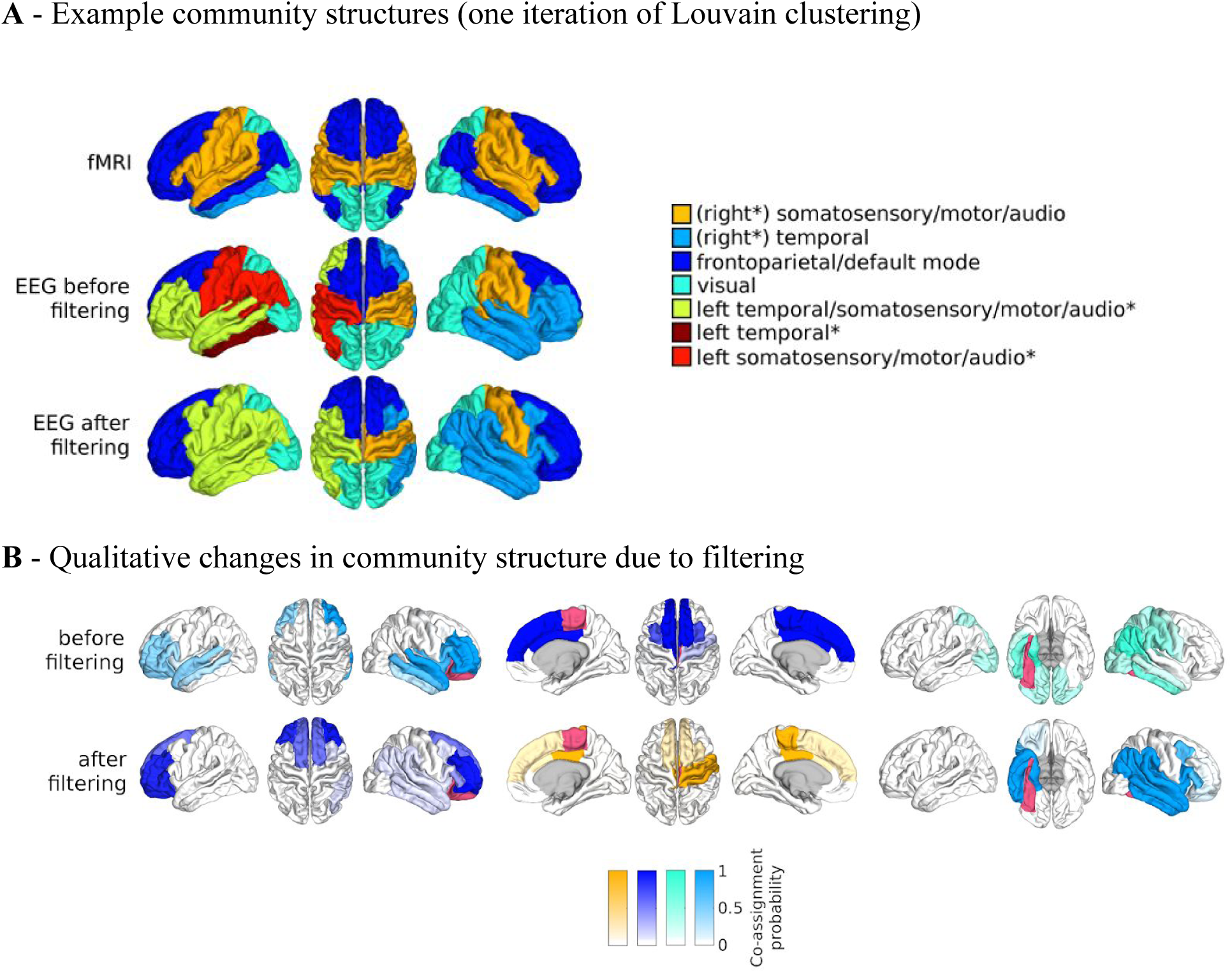
**A**: One result of Louvain clustering community assignments. For EEG, ɣ=1.1, G=400 for the filtered version. In the legend, “*” marks communities only present in EEG. **B**: Surface renderings of the rows/columns of the matrices shown in Figure 7 corresponding to three example ROIs that tend to switch community membership due to filtering. Colors reflect the ROIs’ network membership according to the example in panel A. Left: right lateral orbitofrontal ROI; middle: right paracentral lobule; right: left fusiform area.

We quantify the similarity between the community structures in EEG- and fMRI-FCs by taking the rank correlation between each row/column of the “community matrices” (examples for ɣ=1.1 in Figure 7). We compute this measure depending on two parameters: 1) the resolution parameter ɣ which controls the spatial resolution of the Louvain algorithm and thus the number of communities, and 2) the filter weight G. Figure 9A shows the averaged (over ROIs) similarity in community structure. There is a region where the average rank correlation is ~0.45, i.e. for filter weights between 200 and 1000 and ɣ≥1.075. At the same time, Figure 9B shows that with increasing ɣ, the number of communities also increases: The overall maximum in average rank correlation is 0.60 at ɣ=1.3 and G=800 (Figure S14), but at this point, we have 27 communities. Due to the coarseness of the parcellation used here, and according to the literature (Yeo et al., 2011), we seek to partition the cortex into as few communities as possible while also achieving a good agreement between the community structures of EEG- and fMRI-FCs. Choosing ɣ=1.1 (row indicated in Figures 9A and B), the number of communities is 5 and the agreement with the fMRI community structure is 0.45 (at G=200, 300, or 400, see Figure 9C). The same fit can be achieved at ɣ=1.125 and G=300, but at this point we have 6.5 communities on average. Similarly, without filtering, the correlation is 0.42, but the number of communities is 6. Thus, ɣ=1.1 and G=200, 300, or 400 represents the optimal tradeoff between number of communities and community agreement in comparison to other filter weights, including G=0.

**Figure 9:**
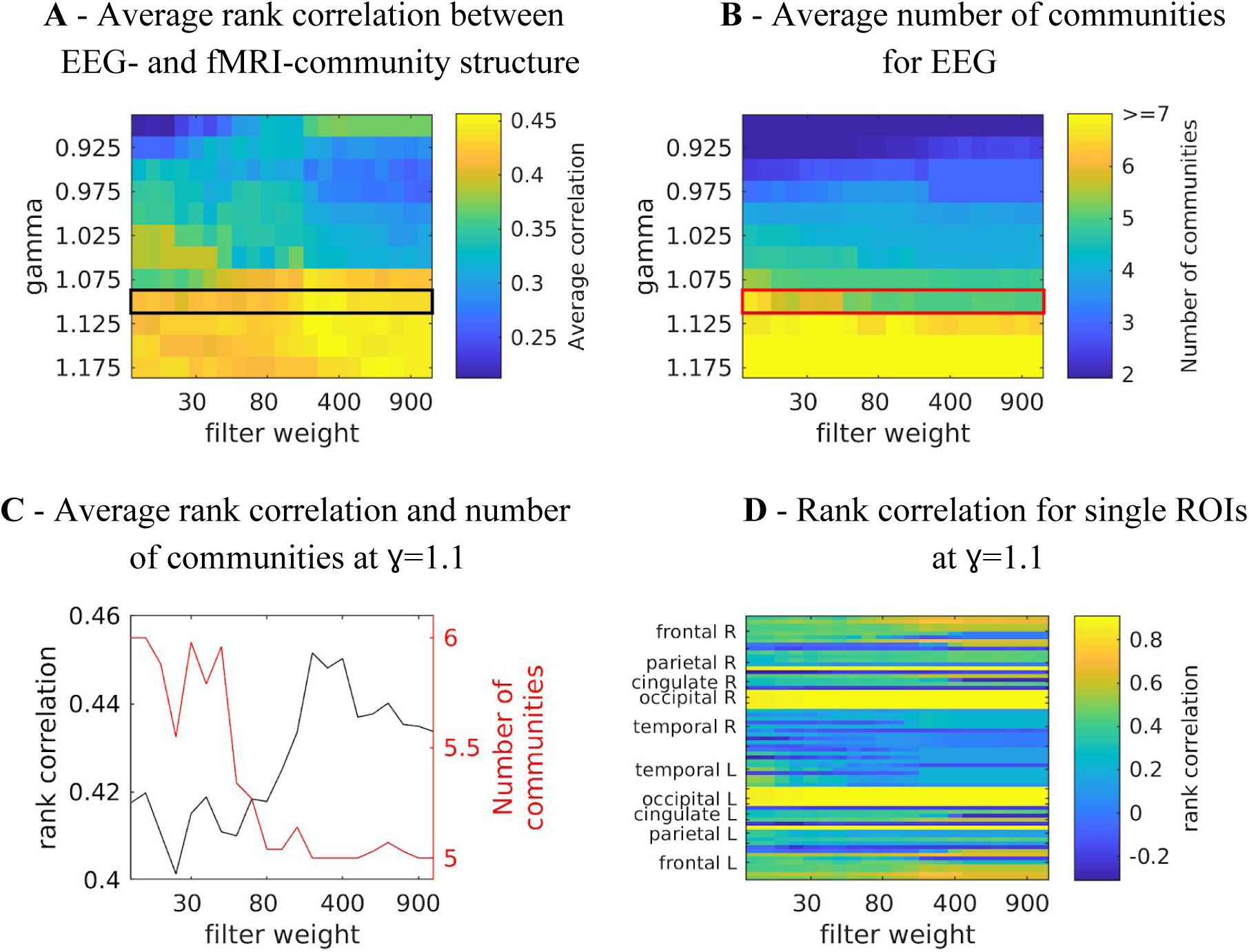
**A**: Agreement between community structure of EEG- and fMRI-FC as measured by average rank correlations between “community matrices” (Figure 7). The black box marks the ɣ which is shown in panel C. **B**: Average number of communities found by the Louvain clustering algorithm. The red box marks the ɣ which is shown in panel C. **C:** Rank correlations and number of communities for ɣ=1.1 (marked in the same colors in panels A and B). **D**: Rank correlations between rows/columns of “community matrices” (Figure 7) of EEG- and fMRI-FC, for ɣ=1.1 (marked with colored boxes in panels A and B).

Since Figure 8A only provides an example partition, we plot on the cortex rows/columns of the community matrices in Figure 7 in order to analyze the community structure across all iterations of the clustering, i.e., the probability of a specific ROI to be co-assigned to the same community as each of the other ROIs. We call this the “community behavior” of this ROI. In Figure 8B, we show three ROIs that exhibit a switch in community membership pertaining to three major communities found by the Louvain algorithm: the frontal community, the somatosensory/motor network, and one of the two temporal communities; we do not show the occipital network as it is quite stable.

Figure 8B, left panel, shows how the “community behavior” of an example frontal ROI (right lateral orbitofrontal ROI) differs between unfiltered and filtered EEG-FCs. The most conspicuous change that is introduced by the filtering is the establishment of a coherent frontal network which includes orbitofrontal regions, as in fMRI. This can be observed both in Figure 8A (blue network) as well as examining the changes in single ROIs displayed in Figure 9D. Figure 8B, middle panel, shows an example from the somatosensory/motor network (right paracentral lobule). This ROI shows a marked improvement in its agreement with fMRI. Namely, before filtering, this ROI is grouped with frontal regions (dark blue network in Figure 8A). Afterwards, it becomes a member of the somatosensory/motor network (orange network in Figure 8A), as is the case in fMRI. However, in contrast to fMRI, a symmetrical network is not established. When using the SC graph for the smoothing procedure, this network is robustly expressed in a certain area of the parameter space (Figures S15 and S16), however, the overall correspondence between EEG- and fMRI-FCs and community structures is much lower at this point (average rank correlation: 0.22).

While the visual network is well-established even without any filtering (Figure 8A, cyan network, and Figure 9D), temporal regions are mostly grouped due to their anatomical proximity. Figure 8B, right panel, shows how the community behavior of an example temporal region (left fusiform area) changes due to filtering. Despite there being a qualitative improvement in that the region switches to a temporal network and decouples from visual regions, there are also many spurious co-assignments with other temporal regions which, in total, explain why this region shows no improvement due to filtering.

## Discussion

In this study, we show that additional information in the SC can be used to selectively increase the FC between brain regions that are connected via white matter fibers, leading to a large-scale functional network structure that is more in accordance with canonical resting state networks (Liu et al., 2017; Yeo et al., 2011). We first show that the strength of FC between pairs of regions that are connected via SC is higher than between those pairs that are not, confirming the assumption that both volume conduction and SC contribute to Euclidean distance-dependent functional connectivity. Second, we find that using a graph that combines SC and Euclidean distances to smooth functional EEG signals in nearest neighbor-graph space results in a higher agreement between fMRI- and EEG-FC structure.

### SC is outperformed by graphs (partially) derived from Euclidean distance

The result that a combination of SC and Euclidean distance performs best was contrary to our initial expectation that SC (fiber counts) would give the largest improvement due to strengthening of FC between distant pairs of ROIs. SC is correlated with Euclidean distance with r=0.48, and there exist many pairs of regions that are *both* nearby in terms of Euclidean distance *and* strongly connected according to the SC (Figure S2A). Thus, strong connections exist between these pairs in all four versions of the graph, and filtering based on any of the four graphs increases the strength of these connections (Figure S4). The increase in fit to the fMRI-FC common to all graphs indicates that despite the fact that these connections are already strong in the EEG-FCs, they are still underestimated compared to fMRI-FC. This is consistent with our rationale that strengthening these connections corresponds to boosting genuine FC mediated by white matter connections.

Furthermore, the GLM analysis showed that the 3rd-strongest predictor for fMRI-FC was an interaction between SC and Euclidean distance, explaining why, beyond the commonalities across graphs based on Euclidean distance, the graph containing information from both SC and Euclidean distances outperforms all other graphs.

A trivial explanation for the increase in agreement between EEG- and fMRI-FC is that graph filtering removes pairwise non-linear relationships which are only present in EEG but not in fMRI. Indeed, Messé et al. (2015) showed that a generative model of fMRI-FC which takes into account only linear relationships between structurally connected pairs outperformed all other models which included non-linearities. In order to investigate this possibility, we simulated white Gaussian noise (WGN) and applied the same filtering procedure, obtaining WGN-FC matrices. We found that at G=100, where the best fit (r=0.51) between EEG-FCs and fMRI-FC is reached, the fit of WGN-FCs to fMRI-FC is 0.35 (Figure S6A), despite the EEG-FC being quite similar to filtered WGN-FCs (average maximum correlation: 0.69, Figure S6B). This suggests that making EEG-FCs more similar to linearly related WGN cannot fully account for the increase in fit between EEG- and fMRI-FC. However, at high G-values (G>500), the WGN-FCs did show a fit to the fMRI-FC that was just as good as that obtained through our filtering procedure with the empirical EEG-FCs, namely around 0.5, which is also the correlation between fMRI-FC and Euclidean distance. At this point, the correlation between WGN-FCs and EEG-FC reaches ~0.9. This indicates that indeed, imposing an interesting graph structure on white Gaussian noise is able to account for about 25% of the variability in fMRI-FC. This result is in line with Messé et al. (2015). Importantly though, our filtering procedure identifies a range of G where only about half of this variance is explained by WGN, suggesting that some interesting non-linearities are preserved.

It is worth mentioning that the results of both ED match and SC might be improved by optimizing the thresholding procedure. As there is currently no consensus on how thresholding of fiber count and recurrence of pairwise connections across subjects should be combined, how the differences between long and short fibers should be taken into account, and how thresholding procedures should differ depending on scanning protocols (e.g. DTI vs DSI), fiber tracking algorithms (e.g. probabilistic vs deterministic), and number of subjects, a systematic exploration of the effect of thresholding procedure is beyond the scope of this study. Nevertheless, the fitting procedure between EEG- and fMRI-FC described here could be employed as one measure to evaluate such thresholding procedures, as a higher fit could be taken to mean that the most relevant fibers were correctly preserved.

### Comparison to methods that attenuate volume conduction

We used envelope-based correlations which are known to be strongly influenced by volume conduction. We opted for this measure because it is widely used (Cabral et al., 2014; Hipp et al., 2012; O’Neill et al., 2015) and captures predominantly slow power modulations, which is closer to what the BOLD signal captures. Also, the graph filtering does not remove zero-lag correlations.

We repeated our analyses using coherence and imaginary part of coherence (Figures S8 and S9), the latter of which is thought to remove zero-lag correlations and thus, volume conduction (Nolte et al., 2004). We found that the increase in fit to the fMRI-FC was much smaller. However, this was the case for both measures, and may therefore just reflect the fact that they are less suited for comparison with BOLD-FC. This does not mean that graph filtering does not work for coherence-based measures, but that comparing to fMRI-FC is not suitable in this case. Further research is necessary in order to clarify the effect of FC measure on graph filtering results. Qualitatively, the results were the same with all three measures.

We also used the method described in Colclough et al. (2015) to orthogonalize the EEG signals, and found that this resulted in a decrease in the correlations between EEG-FCs and fMRI-FC for every subject, suggesting that orthogonalization removes genuine FC.

Overall, these results suggest that both orthogonalization and imaginary coherence remove zero-lag correlations relevant to large-scale network structure. This does not mean that these methods are not useful in detecting true connectivity (Nolte et al., 2004)Some recent studies using such measures have shown network structure partly concordant with fMRI resting state networks, and have added some directionality in network analysis (Coito et al., 2016, 2019; Silfverhuth et al., 2012).

### Euclidean distance and fiber count differentially affect FC in different communities

Our community analysis, using Louvain clustering, revealed that EEG functional networks are differentially affected by our filtering procedure (using “ED match”). For frontal regions, robust improvements are observed, yielding a network that resembles that found in fMRI, apart from missing functional connections that would constitute the hallmarks of the default mode network, i.e. long-range connections between frontal regions and the middle temporal, inferior parietal, and posterior cingulate cortices. However, note that with this coarse anatomical - not functional - parcellation, even the fMRI community structure does not clearly resolve the DMN, which is mixed with the frontoparietal network. This indicates that for this network, “ED match” is a good choice.

For temporal regions, fMRI shows a distinct structure which is not reproduced by EEG either with or without filtering. The main difference is that there is no FC across hemispheres between temporal regions. This shows a limitation of our approach (see below for more discussion on limitations), because even in the SC, there are very few, if any, white matter fiber tracts between the temporal lobes. This is because these fibers are very long and pass through the corpus callosum, making them hard to track. Beyond that, in fMRI, superior, middle, and inferior temporal gyri belong to different networks (somatosensory/motor/auditory, default mode/frontoparietal, limbic/visual, respectively – the resolution of this parcellation is too coarse to resolve these systems properly), while in EEG, the anatomical architecture of the lobes determines the partition into communities. This is a shortcoming which could potentially be improved by using a more EEG-appropriate parcellation.

Finally, for parietal regions typically belonging to a prominent somatosensory/motor network, improvements are achieved using “ED dens”, however, the typical symmetric network including pre- and postcentral gyri is not established. This is due to the fact that the pre- and postcentral gyri are elongated and therefore, the Euclidean distances between ROI centers - which was used to establish the weights in the “ED match” graph - is quite high. Therefore, for this network, SC is the better choice even though overall, SC is outperformed by all ED-based graphs. These observations are in line with recent findings that show that the alignment between SC and FC, i.e. the degree to which FC is shaped by SC, differs across regions (Preti & Van De Ville, 2019), with sensory regions - i.e. the visual and somatosensory/motor cortices - being more strongly aligned with the SC than higher cognitive areas. It may also correspond to a cortical gradient showing different levels of local recurrent connections across functional networks (Wang et al., 2019).

Previous studies have shown consistent functional connectivity between certain homotopic regions (Hipp et al., 2012; Mehrkanoon et al., 2014) while here and elsewhere (Cabral et al., 2014) these connections are shown to be underestimated in M/EEG-FC. On the one hand, these findings might depend on the exact methodology (e.g. which source reconstruction algorithm is used, whether signals were orthogonalized or not). On the other hand, our finding that frontal/occipital networks are quite symmetric across hemispheres even before filtering while temporal and parietal networks are not suggests that FC might be robust between some regions and largely absent between others.

### Limitations

We have already mentioned two limitations of our methodology. First, the SC has obvious shortcomings like the absence of many interhemispheric fibers and the underestimation of long fibers, which are hard to track (Jeurissen et al., 2019; Jones, 2010). Second, we used envelope-based correlations which are known to be strongly influenced by volume conduction. Although we compared to coherence and imaginary part of coherence, a closer investigation of how graph filtering impacts different measures is warranted.

Additionally, the standard FreeSurfer parcellation is probably not optimal for EEG. The ROIs of this parcellation are mostly anatomically defined, not taking into account the nature of the EEG signal: Regions are highly unequal in size, resulting in a wide range of numbers of dipoles being averaged to obtain the ROI time courses. Furthermore, ROIs are in many cases elongated, while for EEG, more spherical regions (as far as this would be anatomically/functionally plausible) would be preferable. Finally, the appropriate number of ROIs is a matter of debate, as a simple correspondence between the number of ROIs and the number electrodes is not applicable (Farahibozorg et al., 2018). This is also in line with our finding that in the GLM, the relative regional variance (RRV) is the second-strongest predictor of EEG-FC in terms of added explained variance, indicating that noise is unequally distributed across ROIs. This could be due to the fact that signals from deep sources are harder to pick up than those of superficial sources (Whittingstall et al., 2003)

On the conceptual level, it is unclear how much EEG-FC should resemble fMRI-FC, since BOLD and the EEG signal are related in a way that is not straightforward. At the same time, recent studies show that RSNs are quite similar across these modalities and therefore, on this level of resolution and detail, we should expect a good agreement between the FC matrices (Coito et al., 2019; Liu et al., 2018). Still, validation should include a biophysical model which simulates both genuine FC based on SC as well as volume conduction.

### Conclusions and future work

Taken together, we add to the thus far sparse knowledge on how SC and FC are related in EEG source space. Developing these methods is crucial for taking full advantage of the immense richness of the EEG signal in the temporal and frequency domain, and combining EEG with other modalities like MEG and fMRI. We have limited our analysis to the grand-average FC, but our method could be used to improve SNR on the single trial level, potentially easing statistical analysis of task-EEG in source space. One important conclusion from our results is that EEG-specific parcellation schemes are necessary to guarantee that we take full advantage of the richness of the EEG signal. Furthermore, our results confirm that it is feasible and sensible to use dynamical models that assume functional activity to spread through white matter fibers in EEG (Bhattacharya et al., 2011; de Haan et al., 2012; Finger et al., 2016; Pons et al., 2010; Ponten et al., 2010; van Dellen et al., 2013). On the data analysis side, this study provides a justification and an avenue to applying more sophisticated methods like graph signal processing (Shuman et al., 2012).

## Methods

### EEG Data and source projection

Data were recorded from 21 healthy controls as part of an epilepsy study at the EEG and Epilepsy Unit, University Hospital of Geneva. This study was approved by the local ethics committee. 3 subjects were excluded due to too many movement artifacts, leaving 18 for analysis. Since subjects were age-matched to patients (not analyzed here), 6 subjects aged less than 18 years were included (age range: 8 to 54 years, median: 29.5). Since we could not find any qualitative differences when excluding these subjects, we proceeded with using all 18 available datasets.

Resting state EEG was collected with the Geodesic Sensor Net with 256 electrodes (Electrical Geodesic, Inc., Eugene, USA) during resting state. Electrodes on cheeks and neck were excluded, leaving 204 electrodes for analysis. Data were downsampled to 1 kHz and artifacts were removed by Infomax-based ICA prior to source projection. Remaining artifacts were marked manually and visually and markers were later used to extract artifact-free intervals of varying length and number per subject (Table S3). Inverse solutions were computed using LAURA with LSMAC as implemented in CARTOOL (Brunet et al., 2011), employing individual head models that were extracted from T1-weighted images (acquired as magnetization prepared rapid-gradient echo MPRAGE volumes with a Siemens TrioTim 3T MRI scanner and a tfl3d1ns pulse sequence with flip angle = 9°; echo time = 2.66ms, repetition time = 1.51s, inversion time = 0.9, voxel size = 1×1×1mm^3^ head first supine) obtained from the same subjects in order to create the forward model. Segmentation and ROI extraction (i.e., parcellation) was performed using Connectome mapper 3 (Tourbier et al., 2020). Gray and white matter were segmented from the MPRAGE volume using Freesurfer with the Lausanne 2008 multiscale parcellation (Hagmann et al., 2008), whose first scale corresponds to the Desikan atlas (Desikan et al., 2006).

Data were source projected to ~5000 dipole locations equally spaced on a 3-dimensional grid, where the gray matter volume extracted from the same images served as a constraint for the dipole locations. In order to project the 3-dimensional time courses of the solution points to 1-dimensional ROI time courses for further analysis, the main direction of variance was extracted using singular value decomposition (Rubega et al., 2019): all solution points were concatenated and their time courses were projected onto the first principal component, preserving most of the variance. This was done for each artifact-free interval (Table S3). Note that all analysis steps were done on the individual level. Since both functional (see below) and structural connectivity were computed between ROIs, averaging of the corresponding FC and SC matrices is possible without ever co-registering images to a common space.

### MRI data

We used three separate structural connectivity datasets in this study, one primary and two additional datasets to check for replicability. See Figure S12 for a comparison between the average SC matrices resulting from these datasets.

Primary structural connectivity matrices and fMRI functional connectivity matrices: 88 healthy control subjects (mean age 29.7 years, minimum 18.5, maximum 59.2 years; 34 females) were scanned in a 3-Tesla MRI scanner (Trio, Siemens Medical, Germany) using a 32-channel head-coil. Informed written consent in accordance with institutional guidelines (protocol approved by the Ethics Committee of Clinical Research of the Faculty of Biology and Medicine, University of Lausanne, Switzerland, #82/14, #382/11, #26.4.2005) was obtained for all subjects. A large subset (70 out of 88 subjects) of the structural and functional connectivity matrices obtained from this data are available on Zenodo (Griffa et al., 2019), where the processing pipelines are described in detail. Briefly, for diffusion imaging, a DSI sequence (128 diffusion-weighted volumes and a single b0 volume, maximum b-value 8,000 s/mm^2^, 2.2×2.2×3.0 mm voxel size) was applied. For resting state fMRI, a gradient echo EPI sequence sensitive to BOLD contrast (3.3-mm in-plane resolution and slice thickness with a 0.3-mm gap, TR 1,920 ms, resulting in 280 images per participant) was applied. A magnetization-prepared rapid acquisition gradient echo (MPRAGE) sequence sensitive to white/gray matter contrast (1-mm in-plane resolution, 1.2-mm slice thickness) was also acquired. ROI extraction (i.e., parcellation) was performed using the Connectome mapper (Daducci et al., 2012). Gray and white matter were segmented from the MPRAGE volume using Freesurfer with the Lausanne 2008 multiscale parcellation (Hagmann et al., 2008), whose first scale corresponds to the Desikan atlas (Desikan et al., 2006). DSI data were reconstructed following the protocol described in (Wedeen et al., 2005).

Structural connectivity matrices were estimated for individual participants using deterministic streamline tractography on reconstructed DSI data, initiating 32 streamline propagations per diffusion direction, per white matter voxel (Wedeen et al., 2008).

FMRI volumes were corrected for physiological variables, including regression of white matter, cerebrospinal fluid, as well as motion (three translations and three rotations, estimated by rigid body co-registration). Time courses of voxels falling into each ROI were average and FC was computed as correlations between these ROI time courses.

Control structural connectivity matrices - small DTI cohort: 20 healthy control subjects (17 females, mean age: 23, age range: 20-29 years) were scanned in a 3-Tesla MRI scanner (DISCOVERY MR750, GE Healthcare, USA) using a 32-channel head-coil. For diffusion imaging, a DTI sequence (30 diffusion-weighted directions with b-value 1000s/mm^2^ and 5 b0 volumes, with 1×1×2.2 mm^3^ voxel size, TE/TR=87/8000 ms) was applied. A T1-weighted (T1w) volume (1mm isotropic resolution) was also acquired, using an inversion-recovery-prepared fast spoiled gradient recalled brainvolume (IR-FSPGR-BRAVO) sequence, a fast SPGR sequence with parameters tuned to optimize brain tissue contrast. ROI extraction (i.e., parcellation),, diffusion signal analysis, and construction of SCs were performed using the Connectome mapper 3 (Tourbier et al., 2020). Gray and white matter were segmented from the T1w volume using Freesurfer and the Lausanne 2008 multiscale parcellation scheme (Hagmann et al., 2008). DTI data was corrected for motion and distortions using FSL mcflirt (Jenkinson et al., 2002)and eddy_correct (Andersson & Sotiropoulos, 2016), from which fiber orientation distribution functions (fODFs) were estimated via constrained spherical deconvolution (order 4). Then, by seeding from the white matter voxels and by the means of fODFs-driven deterministic tractography (MRtrix3 SD_STREAM, Tournier et al., 2019), a total of 1M streamlines were reconstructed. T1w and b0 mean volumes were non-linearly co-registered using ANTs (Avants et al., 2008) symmetric normalization in order to project the extracted ROIs to the DTI data space. Finally, by identification of the ROIs connected by fibers (Hagmann et al., 2008), a graph is constructed and the structural connectivity matrices are estimated for each individual.

Control structural connectivity matrices - Human Connectome Project (HCP) Retest cohort (Feinberg et al., 2010; Moeller et al., 2010; Setsompop et al., 2012; Van Essen et al., 2012): The Human Connectome Project (HCP) Retest cohort consists of a subset of 45 HCP subjects recruited by the HCP to undergo the HCP 3T scanning pipeline a second time. The HCP uses a customized Siemens 3T “Skyra Connectome” scanner with a 32-channel head-coil, which is located at Washington University in St Louis, USA, and which provides an increased gradient strength compared to standard scanners. This allows the acquisition of multi-shell dMRI data (voxel size: 1.25×1.25×1.25 mm3). 90 diffusion weighting directions and 6 b0 volumes were acquired for three different shells (with b=1000, 2000, and 3000 s/mm^2^, voxel size: 1.25×1.25×1.25 mm^3^; Feinberg et al., 2010). An MPRAGE image is also acquired sensitive to the white/gray matter contrast (0.7-mm isotropic resolution) and used for ROI extraction. In this work we used in particular the publicly available T1w and dMRI derived data preprocessed by the “HCP minimal preprocessing pipelines” described in Glasser et al. (2013). This consists of the (1) T1w images, (2) the corresponding processed Freesurfer outputs, and (3) the fully corrected dMRI images. ROI extraction (i.e., parcellation), diffusion signal reconstruction and tractography, and construction of SCs were also performed using the Connectome mapper 3 (Tourbier et al., 2020). Gray and white matter were segmented using the existing pre-computed Freesurfer outputs and the Lausanne 2008 multiscale parcellation scheme. Fiber orientation distribution functions (fODFs) were estimated from the existing preprocessed dMRI volumes via constrained spherical deconvolution (order 10). Then, by the means of anatomically-constrained and fODFs-driven probabilistic tractography (MRtrix3 ACT iFOD2, ACT [Smith et al., 2012]) with seeding from the white matter interface, a total of around 200K streamlines were reconstructed. T1w and b0 mean volumes are non-linearly co-registered using ANTs symmetric normalization and the extracted ROIs are projected to the dMRI data space. Finally, by identification of the ROIs connected by fibers, a graph is constructed and the structural connectivity matrices are estimated for each individual.

### Functional connectivity measures for EEG

For our main analysis, we used power envelope correlations to measure functional connectivity in alpha (8-13 Hz), beta (13-30 Hz), and gamma (30-40 Hz) bands. All data analysis was performed in Matlab 2017a (MathWorks, Natick, USA). We extracted artifact-free intervals from the resting state time courses by using the artifact markers mentioned above, admitting only intervals of at least 19s duration in order to guarantee a reliable estimation of slow envelope modulations (see below). Prior to extracting the FCs, for each artifact-free interval, the data were downsampled to 250 Hz in order to avoid numerical errors which can be induced if the band of interest is much lower than the sampling rate.

Envelope correlations were computed as in Cabral et al. (2014): after zero-phase band-pass filtering (eegfilt from EEGLAB software package) the EEG single trial time series in the respective bands, envelopes were extracted using the Hilbert transform. Envelopes were again low-pass filtered at 0.5 Hz, capturing ultraslow modulations in the respective bands. This measure was chosen to be as closely related as possible to fMRI-FC (Pearson correlation between BOLD time courses).

As a comparison (see Discussion), we computed the EEG-/fMRI-FC-correlations also using FCs obtained with coherence and imaginary part of coherence (icoherence, Nolte et al., 2004). This part of the analysis was performed using functions implemented in the Brainstorm toolbox (Tadel et al., 2011), which is documented and freely available for download online under the GNU general public license (http://neuroimage.usc.edu/brainstorm). Brainstorm estimates (i)coherence in a minimum number of 5 windows which overlap by 50%. At a sampling rate of 250 Hz, this results in a minimum required signal length of ~3s. Thus, artifact-free intervals were segmented into epochs of ~3s. Since we found that these measures were more susceptible to noise introduced by remaining artifacts than envelope correlations, we further rejected epochs which, at any point, deviated from the mean by more than 6 standard deviations. See Table S3 for the number of epochs that was used for each subject.

### Average structural connectivity matrices

All data analysis was performed in Matlab 2017a (MathWorks, Natick, USA). We used the method introduced in Betzel et al. (2019) to obtain unbiased group-consensus SCs (average number of fibers) for all three datasets used in this study (primary, HCP, and DTI, see above). The reason for using a group average rather than individual SC matrices is that fiber tracking algorithms are not sufficiently reliable on an individual level, and information from the entire group is necessary to identify the most likely true-positive connections.

In brief, this method takes into account the fact that interhemispheric connections are less reliable than intrahemispheric ones. Using a single threshold on the average SC strength (i.e. setting all connections that have less than a certain average number of fibers to 0) results in an underestimated interhemispheric connection density compared to single subjects. The method used here (Betzel et al., 2019) preserves both intra- and interhemispheric connection density found in single subjects by applying separate thresholds. Additionally, we require a recurrence of at least 30%. The resulting connection density in our primary SC is 25% (39% within and 12% across hemispheres).

Applying the distance-dependent threshold (Betzel et al., 2019) and a recurrence threshold of 0.3 to the additional datasets, DTI and HCP as described above, resulted in different connection densities (DTI: 22% overall, 36% within and 8% across hemispheres; HCP: 50% overall, 67% within and 34% across hemispheres). We adjusted the parameters of the distance-dependent thresholding procedure such that the resulting average SC exhibited a density that, for the DTI dataset, was above the density of the single subjects (18% higher), and for the HCP dataset such that it was below the single subject (41% lower). In order to still be able to take advantage of the superior quality of the HCP data compared to the DTI data, we chose to match the intrahemispheric connection density, allowing the interhemispheric, and thus, the overall density, to vary. This resulted, for DTI, in an overall density of 24% (39% within and 9% across hemispheres), and for HCP, in an overall density of 30% (39% within and 20% across hemispheres).

Lastly, the randomized SC used in one of our control analyses was obtained using the Brain Connectivity Toolbox (Rubinov & Sporns, 2010)function randmio_und_connected() with 100 iterations, which we used to randomly rewire the primary SC 100 times, thereby preserving the degree sequence.

### Search information

All data analysis was performed in Matlab 2017a (MathWorks, Natick, USA). In order to correlate FC and SC matrices, we have to overcome the problem that FCs are dense (Pearson correlation of time courses can be computed for all pairs of ROIs), while fiber tracts only exist between a subset of ROI pairs. Here, we use search information as defined in Goñi et al. (2014) to derive a dense matrix SC_SI_ from the SC. A path π_*s→t*_ is a sequence of edges (non-zero entries in the SC) leading from node s to node t. The search information *S*(π_*s→t*_) is then computed as

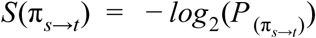

where

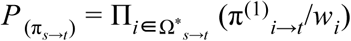

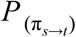 is the probability that a signal travelling from node s to node t will take the shortest path π_*s→t*_. Ω*_*s→t*_ is the sequence of nodes on the shortest path, π^(1)^_*i→t*_ is the first element (edge) on the path from node i to node t, and *w*_*i*_ is the weight of this edge. The intuition is that if there exist many such sequences between two given nodes s and t, the shortest path is “hidden” and more “information” is needed to find it. Figure S1 shows this matrix.

### General linear model

We use Matlab’s function stepwiseglm() to compute coefficients and p-values for a general linear model (GLM) that computes FC from four predictors:

1. search information derived from SC, as detailed above
2. Euclidean distance
3. relative regional variance (RRV)
4. ROI size

RRV serves as a proxy for signal to noise ratio. First, we compute for each ROI and each subject the average (over intervals) variance of this ROI’s time courses. The variances are then scaled such that the maximum (over ROIs) variance is 1 and averaged over subjects.

Matlab’s stepwiseglm() finds the order of predictor variables in terms of their deviances, i.e. twice the difference between the log-likelihood of that model and the full model (using all possible predictors, i.e. 4 main effects and 6 interaction terms). A variable is added into the model if the difference between deviances obtained by adding it is significant (χ2 test). Removal of a term is also possible; this can occur if predictors are linearly dependent. Any non-significant predictors are not included. Interactions are only included if the main effects are significant. For comparison, we fit single-variable GLMs that use only the intercept and one predictor variable at a time (see Figure 2A).

### Smoothing in graph space

Figure 1 illustrates our filtering approach. We performed spatial low-pass filtering, or smoothing, using the space defined by different versions of the structural connectivity graph. Each node of this weighted and undirected graph is one region of interest defined by the Lausanne2008 parcellation scheme (Hagmann et al., 2008), yielding 68 nodes. The edges of the graph are defined by the density of white matter tracts discovered between these ROIs via dMRI. Smoothing of node *i*’s signal at time *t*, *x*_*i*_(*t*), is then performed using the nearest neighbors of node *i*, i.e. all nodes *j* ≠ *i* which have a direct connection to *i* according to the connectivity matrix *C* with entries *c*_*ij*_:

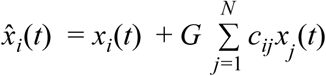

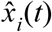 is the resulting filtered signal. *G* is a scalar parameter 0≤*G*≤1000 which tunes the impact of the neighbors’ signal. The filtering procedure was implemented using Matlab 2017a.

### Statistical analysis for comparing EEG- and fMRI-FC matrices

We compute individual FC matrices after smoothing in graph space of each subject’s EEG node time courses. Each subject’s FC matrix, for each value of *G*, was correlated to the group average fMRI-FC matrix. Since the fMRI-FC matrix was computed from a rather large sample (88 subjects), we assume that it represents a “canonical” functional network structure. In order to assess whether the filtering procedure increases each subject’s EEG-FC’s similarity to this canonical network structure, we compared the best fit resulting from each of the four versions of the graph to the baseline as well as among each other, using a Wilcoxon signed-rank test. This is a paired test, comparing the differences of each subject’s individual correlations against 0. Since there are 10 comparisons (each of the four graph versions against baseline, and 7 pairwise comparisons between graph versions), we just show a summary of the results as boxplots, however, note that we did not compare the means or medians, but the individual differences.

### Functional connectivity matrices derived from white Gaussian noise

In order to explore the effect that smoothing in graph space is likely to remove interesting non-linearities in EEG-FC, we created white Gaussian noise (WGN) time series using Matlab’s wgn()-function. We preserved the power (variance) of the original signals for each artifact-free interval separately and computed FC in exactly the same way as for the EEG data. In addition to assessing the correlation between these WGN-FCs and fMRI-FC, we quantified how much filtered EEG-FCs resemble WGN-FCs by computing for each filter weight the correlation between the EEG-FCs filtered with this weight and all filtered WGN-FCs, as a match between the weights cannot be assumed. In Figure S6B, the maximum correlations to any filtered WGN-FC are shown.

### Analysis of community/modular structure of FC matrices

We used the Louvain community detection algorithm implemented in the Brain Connectivity toolbox (Rubinov & Sporns, 2010) to evaluate the community structure present in the FC matrices. This algorithm takes the FC (adjacency) matrix and assigns each ROI to a community. The function takes one parameter, ɣ, which controls the spatial resolution and thus, indirectly, the number of communities that are detected, with higher values of ɣ leading to more and smaller communities. We varied ɣ in a range of 0.9 ≤ γ ≤ 1.3, resulting in a minimum of 2 and a maximum of 35 communities (Figure S14).

Since the community structure depends to some degree on the initial conditions (randomly assigned module memberships of ROIs), we repeated the procedure 200 times. Instead of opting for a “hard assignment” approach, we took into account the uncertainty of the cluster structures. For each round of clustering, we determined for each pair of ROIs whether they were assigned to the same community. This resulted in a “community matrix” which can be interpreted as indicating the probability of a pair of ROIs to belong to the same community. Thus, each row/column of one of these matrices describes the “community profile” of a given ROI. In order to compare the community structures of EEG- vs fMRI-FCs, we compute the rank correlations between the community profiles (row/column of community matrix) of each ROI.

## Acknowledgements

This work was supported by Swiss National Science Foundation Sinergia grant no. 170873 and grant no. 169198 (to S. Vulliemoz).

