## Supplementary Information for "Using structural connectivity to augment community structure in EEG functional connectivity"

Table S1: (related to Figure 2A) Detailed GLM results with outputs of Matlab's stepwiseglm() and explained variances. *Estimate* is the coefficient value, *SE* is the standard error. Only significant effects are listed. For explained variances, the variables are listed in the order in which they were entered into the model; r2 full (cum.) and r2 single corresponds to the explained variance of the predictor variable in the full model, cumulatively, and the GLM in which the variable was the only predictor (additionally to the intercept). SCSI: search information of the structural connectivity, ED: Euclidean distance, RRV: relative regional variance.

### alpha band without filtering

coefficients

| | Estimate | SE | tStat | pValue |
|---|---|---|---|---|
| (Intercept) | 1.24E+00 | 2.02E-02 | 6.14E+01 | 0.00E+00 |
| SCSI | -9.43E-02 | 3.63E-03 | -2.60E+01 | 1.09E-130 |
| ED | -1.19E-02 | 3.09E-04 | -3.84E+01 | 4.49E-249 |
| RRV | 5.05E-01 | 4.32E-02 | 1.17E+01 | 1.11E-30 |
| SC:ED | 1.19E-03 | 4.99E-05 | 2.38E+01 | 5.04E-112 |
| ED:RRV | -4.00E-03 | 6.04E-04 | -6.63E+00 | 4.18E-11 |

explained variances

| | | r2 full (cum.) | r2 single |
|---|---|---|---|
| 1. | ED | 0.55 | 0.55 |
| 2. | RRV | 0.57 | 0.02 |
| 3. | SCSI | 0.59 | 0.14 |
| 4. | SC:ED | 0.67 | 0.36 |
| 5. | ED:RRV | 0.67 | 0.01 |

**beta band without filtering**

coefficients

| | Estimate | SE | tStat | pValue |
|---|---|---|---|---|
| (Intercept) | 1.56E+00 | 3.04E-02 | 5.14E+01 | 0.00E+00 |
| SCSI | -9.78E-02 | 4.67E-03 | -2.09E+01 | 3.41E-89 |
| ED | -1.34E-02 | 4.41E-04 | -3.04E+01 | 3.37E-170 |
| ROIsize | -2.42E-05 | 4.23E-06 | -5.73E+00 | 1.13E-08 |
| RRV | 2.21E-01 | 1.60E-02 | 1.38E+01 | 7.92E-42 |
| SC:ED | 1.07E-03 | 5.51E-05 | 1.95E+01 | 4.63E-78 |
| SC:ROIsize | 3.25E-06 | 6.97E-07 | 4.66E+00 | 3.34E-06 |
| ED:ROIsize | 1.66E-07 | 5.66E-08 | 2.93E+00 | 3.41E-03 |

explained variances

| | | r2 full (cum.) | r2 single |
|---|---|---|---|
| 1. | ED | 0.58 | 0.58 |
| 2. | RRV | 0.59 | 0.01 |
| 3. | SCSI | 0.61 | 0.15 |
| 4. | SC:ED | 0.66 | 0.39 |
| 5. | ROIsize | 0.66 | 0.00 |
| 6. | SC:ROIsize | 0.67 | 0.05 |
| 7. | ED:ROIsize | 0.67 | 0.17 |

**gamma band without filtering**

coefficients

| | Estimate | SE | tStat | pValue |
|---|---|---|---|---|
| (Intercept) | 1.17E+00 | 1.94E-02 | 6.01E+01 | 0.00E+00 |
| SCSI | -8.63E-02 | 3.49E-03 | -2.47E+01 | 1.56E-119 |
| ED | -1.12E-02 | 2.98E-04 | -3.76E+01 | 8.06E-241 |
| RRV | 3.30E-01 | 4.16E-02 | 7.94E+00 | 3.27E-15 |
| SC:ED | 1.06E-03 | 4.80E-05 | 2.22E+01 | 1.28E-98 |
| ED:RRV | -2.11E-03 | 5.81E-04 | -3.63E+00 | 2.87E-04 |

explained variances

| | | r2 full (cum.) | r2 single |
|---|---|---|---|
| 1. | ED | 0.56 | 0.56 |
| 2. | RRV | 0.57 | 0.01 |
| 3. | SCSI | 0.59 | 0.16 |
| 4. | SC:ED | 0.66 | 0.38 |
| 5. | ED:RRV | 0.66 | 0.01 |

**alpha band with filtering** (SC, G=100, best fit to fMRI-FC)
coefficients

| | Estimate | SE | tStat | pValue |
|---|---|---|---|---|
| (Intercept) | 1.78E+00 | 4.14E-02 | 4.30E+01 | 2.30E-296 |
| SCSI | -1.43E-01 | 6.58E-03 | -2.17E+01 | 3.28E-95 |
| ED | -1.48E-02 | 5.24E-04 | -2.83E+01 | 1.53E-151 |
| ROIsize | 1.29E-05 | 5.46E-06 | 2.37E+00 | 1.81E-02 |
| RRV | -2.13E-01 | 7.07E-02 | -3.01E+00 | 2.63E-03 |
| SC:ED | 1.38E-03 | 6.52E-05 | 2.11E+01 | 1.49E-90 |
| SC:ROIsize | -3.42E-06 | 9.44E-07 | -3.62E+00 | 2.97E-04 |
| SC:RRV | 4.75E-02 | 8.69E-03 | 5.47E+00 | 5.03E-08 |
| ED:ROIsize | 4.45E-07 | 6.73E-08 | 6.62E+00 | 4.51E-11 |
| ROIsize:RRV | 2.93E-05 | 9.28E-06 | 3.16E+00 | 1.61E-03 |

explained variances

| | | r2 full (cum.) | r2 single |
|---|---|---|---|
| 1. | ED | 0.41 | 0.41 |
| 2. | SCSI | 0.57 | 0.39 |
| 3. | SCSI:ED | 0.62 | 0.47 |
| 4. | ROIsize | 0.65 | 0.09 |
| 5. | RRV | 0.67 | 0.00 |
| 6. | SCSI:RRV | 0.68 | 0.03 |
| 7. | ED:ROIsize | 0.68 | 0.02 |
| 8. | SCSI:ROIsize | 0.68 | 0.02 |
| 9. | ROIsize:RRV | 0.68 | 8.00E-04 |

**beta band with filtering** (SC, G=100, best fit to fMRI-FC)

coefficients

| | Estimate | SE | tStat | pValue |
|---|---|---|---|---|
| (Intercept) | 1.91E+00 | 4.34E-02 | 4.40E+01 | 1.79E-306 |
| SCSI | -1.33E-01 | 6.90E-03 | -1.93E+01 | 4.72E-77 |
| ED | -1.43E-02 | 5.49E-04 | -2.60E+01 | 1.03E-130 |
| ROIsize | 1.92E-05 | 5.73E-06 | 3.36E+00 | 7.93E-04 |
| RRV | -2.28E-01 | 7.41E-02 | -3.07E+00 | 2.15E-03 |
| SC:ED | 1.21E-03 | 6.83E-05 | 1.78E+01 | 3.04E-66 |
| SC:ROIsize | -2.57E-06 | 9.90E-07 | -2.60E+00 | 9.48E-03 |
| SC:RRV | 4.86E-02 | 9.12E-03 | 5.33E+00 | 1.07E-07 |
| ED:ROIsize | 3.78E-07 | 7.06E-08 | 5.35E+00 | 9.45E-08 |
| ROIsize:RRV | 2.11E-05 | 9.73E-06 | 2.17E+00 | 3.04E-02 |

explained variances

| | | r2 full (cum.) | r2 single |
|---|---|---|---|
| 1. | ED | 0.43 | 0.43 |
| 2. | SCSI | 0.58 | 0.38 |
| 3. | ROIsize | 0.62 | 0.10 |
| 4. | SCSI:ED | 0.66 | 0.49 |
| 5. | RRV | 0.67 | 0.01 |
| 6. | SCSI:RRV | 0.68 | 0.04 |
| 7. | ED:ROIsize | 0.68 | 0.01 |
| 8. | SCSI:ROIsize | 0.68 | 0.01 |
| 9. | ROIsize:RRV | 0.68 | 2.55E-05 |

**gamma band with filtering** (SC, G=100, best fit to fMRI-FC)

coefficients

| | Estimate | SE | tStat | pValue |
|---|---|---|---|---|
| (Intercept) | 1.65E+00 | 3.99E-02 | 4.13E+01 | 7.91E-279 |
| SCSI | -1.35E-01 | 6.34E-03 | -2.13E+01 | 2.96E-92 |
| ED | -1.39E-02 | 5.05E-04 | -2.75E+01 | 2.50E-144 |
| ROIsize | 1.53E-05 | 5.26E-06 | 2.91E+00 | 3.60E-03 |
| RRV | -2.08E-01 | 6.81E-02 | -3.06E+00 | 2.24E-03 |
| SC:ED | 1.28E-03 | 6.28E-05 | 2.03E+01 | 1.19E-84 |
| SC:ROIsize | -2.64E-06 | 9.10E-07 | -2.90E+00 | 3.80E-03 |
| SC:RRV | 4.56E-02 | 8.38E-03 | 5.44E+00 | 5.77E-08 |
| ED:ROIsize | 3.19E-07 | 6.49E-08 | 4.92E+00 | 9.11E-07 |
| ROIsize:RRV | 2.58E-05 | 8.94E-06 | 2.89E+00 | 3.89E-03 |

explained variances

| | | r2 full (cum.) | r2 single |
|---|---|---|---|
| 1. | ED | 0.43 | 0.43 |
| 2. | SCSI | 0.58 | 0.39 |
| 3. | SCSI:ED | 0.63 | 0.48 |
| 4. | ROIsize | 0.66 | 0.08 |
| 5. | RRV | 0.68 | 0.00 |
| 6. | SCSI:RRV | 0.68 | 0.04 |
| 7. | ED:ROIsize | 0.69 | 0.02 |
| 8. | SCSI:ROIsize | 0.69 | 0.03 |
| 9. | ROIsize:RRV | 0.69 | 3.00E-04 |

**fMRI**

coefficients

| | Estimate | SE | tStat | pValue |
|---|---|---|---|---|
| (Intercept) | 6.78E-01 | 2.50E-02 | 2.72E+01 | 5.02E-141 |
| SCSI | -5.58E-02 | 4.26E-03 | -1.31E+01 | 7.69E-38 |
| ED | -4.76E-03 | 3.31E-04 | -1.44E+01 | 5.19E-45 |
| ROIsize | 2.91E-06 | 1.74E-06 | 1.67E+00 | 9.42E-02 |
| RRV | -1.65E-01 | 6.05E-02 | -2.73E+00 | 6.43E-03 |
| SC:ED | 5.35E-04 | 5.27E-05 | 1.02E+01 | 1.03E-23 |
| SC:RRV | 3.02E-02 | 7.26E-03 | 4.15E+00 | 3.41E-05 |
| ED:RRV | -2.37E-03 | 7.26E-04 | -3.27E+00 | 1.11E-03 |
| ROIsize:RRV | 1.58E-05 | 7.67E-06 | 2.06E+00 | 3.94E-02 |

explained variances

| | | r2 full (cum.) | r2 single |
|---|---|---|---|
| 1. | ED | 0.19 | 0.19 |
| 2. | SCSI | 0.23 | 0.14 |
| 3. | SC:ED | 0.27 | 0.18 |
| 4. | RRV | 0.28 | 0.03 |
| 5. | ROIsize | 0.28 | 0.03 |
| 6. | SC:RRV | 0.28 | 0.05 |
| 7. | ED:RRV | 0.29 | 0.08 |
| 8. | ROIsize:RRV | 0.29 | 0.01 |

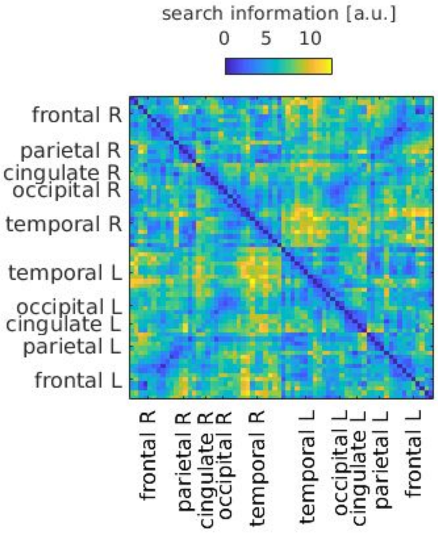


Figure S1: (related to Results - “SC provides additional predictive power for EEG-FC“ and Methods- “Search Information”) Search information matrix used in the GLM and FC-correlation analyses.

**A** - Relationship between SC, FC, and Euclidean distance

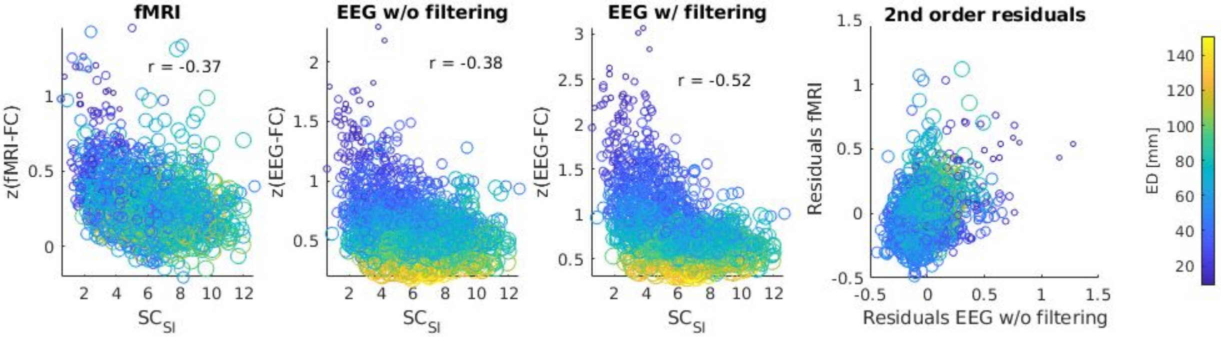


**B** - Distribution of 1st order residuals

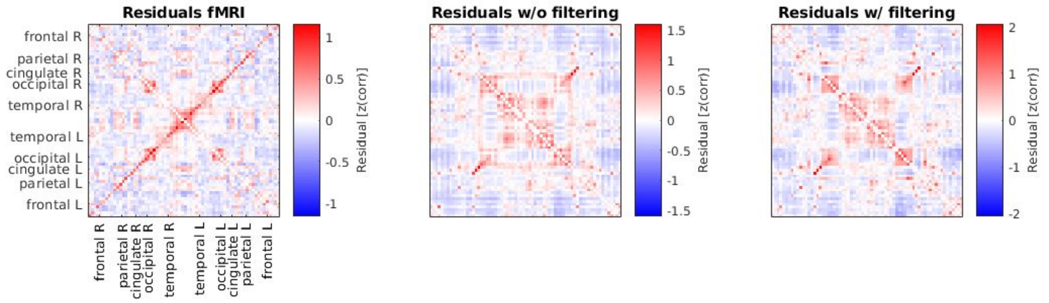


Figure S2: (related to GLM analysis) **A**: In the first three panels, z-transformed FC-values are plotted against $SC_{SI}$ for fMRI, EEG before and EEG after filtering (G=100, ED match). In the forth panel, the FC values that remain unexplained after regressing out Euclidean distance as well as $SC_{SI}$ (2nd order residuals) are plotted for EEG and fMRI in order to highlight the differences between the modalities. A positive residual means that the FC was underestimated, a negative one, overestimated. Sizes and colors of the circles code for the Euclidean distance. **B**: Residuals of single-variable GLM which predicts FC solely from $SC_{SI}$.

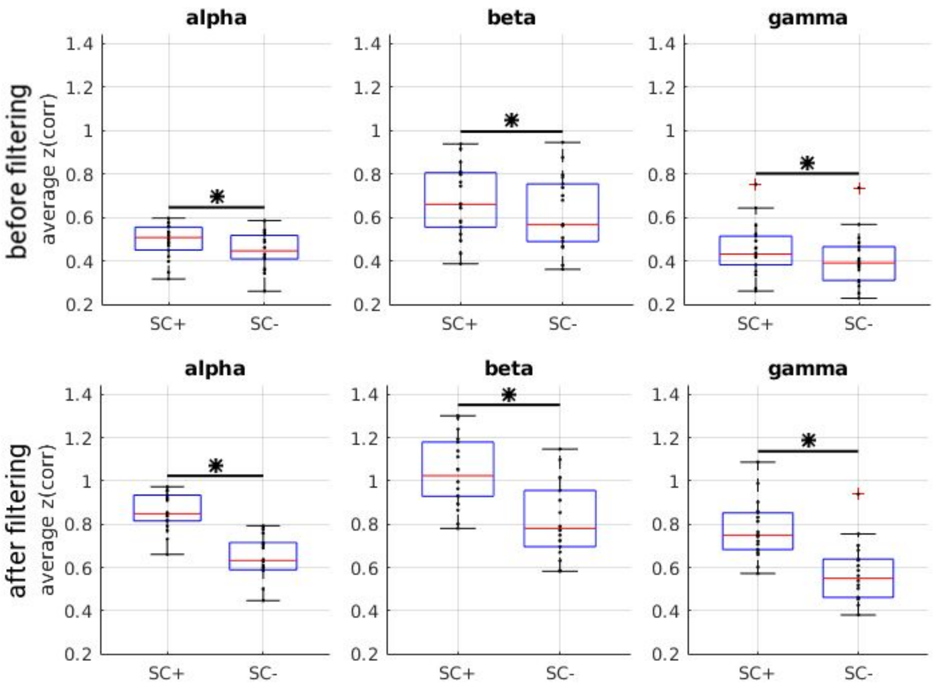


Figure S3: (related to Figure 2B) Comparison between average EEG-FC values for pairs that are connected by SC (“SC+”) and those that are not (“SC-”), before (top row; identical to main manuscript) and after (bottom row) filtering. The samples that are compared are matched in their ED distribution to control for the fact that pairs that are connected tend to be closer together than those that are not. Stars mark significant results according to the Wilcoxon signed-rank test at alpha=0.05 (Bonferroni-corrected for multiple comparisons).

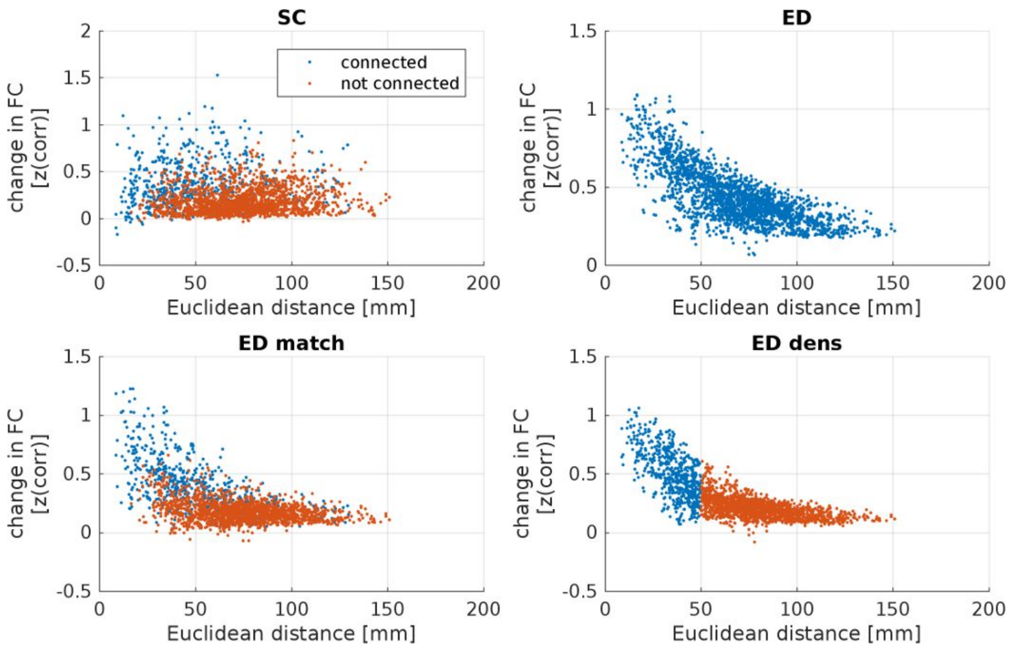


Figure S4: Illustration of how changes in EEG-FC (beta band, G=100) depend on Euclidean distance and on the presence of a direct connection for each of the four matrices used as filters.

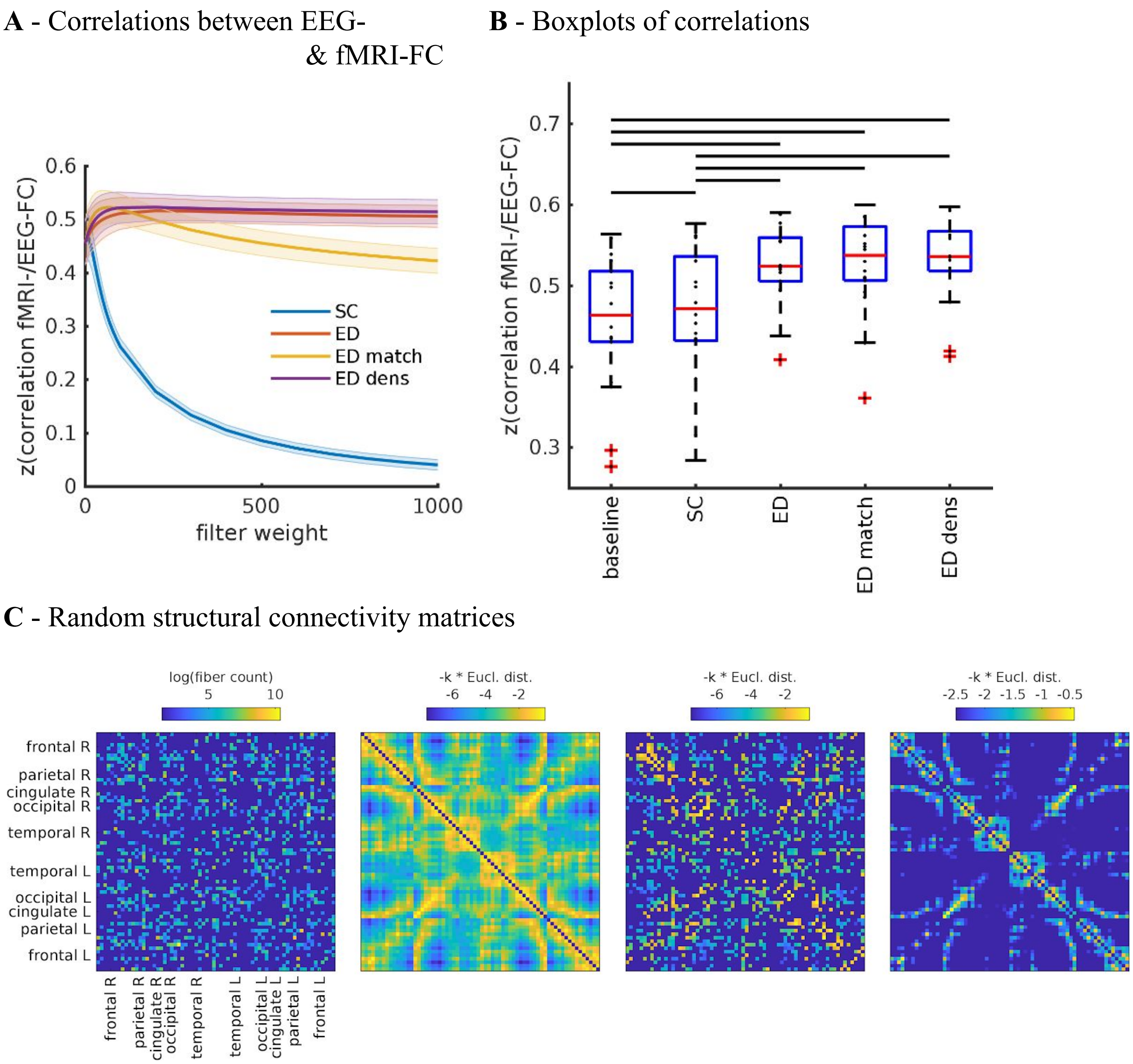


Figure S5: (related to Results - "Graph filtering increases resemblance between EEG-FC and fMRI-FC", Figure 4) **A**: Fit (z-transformed correlation) between EEG-FCs (beta band) obtained using a randomized SC-matrix and the fMRI-FC. The shaded regions mark the 95% confidence interval. **B**: Boxplots summarizing results of the Wilcoxon signed-rank test comparing individuals' maximum fits (shown in panel A) across versions of the SC as well as to the baseline correlation between unfiltered EEG-FCs and fMRI-FC. Black bars mark significant differences. Red lines mark the median, each black dot marks the value for one subject. Note that we did not compare the medians, but the individual differences (see Methods). **C**: All SCs from which graphs for filtering are derived, as in Figure 3 in the main manuscript, and matching panels A and B, i.e. order of the SCs (from left to right) is "SC", "ED", "ED match", "ED dens". Note that the second and forth SC are identical to the original SCs (Figure 3), as the Euclidean distances are the same, and the density is identical.

**A** - Correlation of white Gaussian noise-FC with fMRI-FC

**B** - Correlation of filtered EEG-FCs with white Gaussian noise-FCs

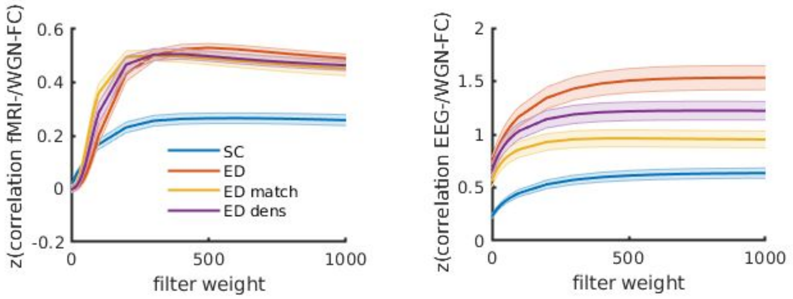


Figure S6: (related to Results) Results of graph filtering applied to white Gaussian noise (WGN). **A**: Correlation of FCs derived from filtered WGN with fMRI-FC (similar to Figure 4A). **B**: Similarity between filtered EEG-FCs and WGN-FCs. For each value of G, the best-fitting WGN-FC (out of all values of G) was selected and the correlation is shown.

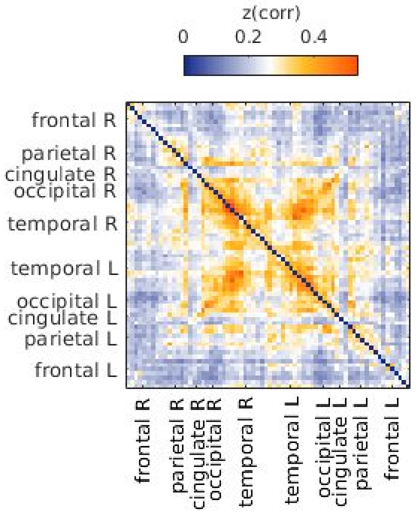


Figure S7: (Related to Results: “Graph filtering increases resemblance between EEG-FC and fMRI-FC“, and Discussion: “Comparison to other methods that attenuate volume conduction”) Average EEG-FC (without filtering) after each individual’s EEG signals were orthogonalized.

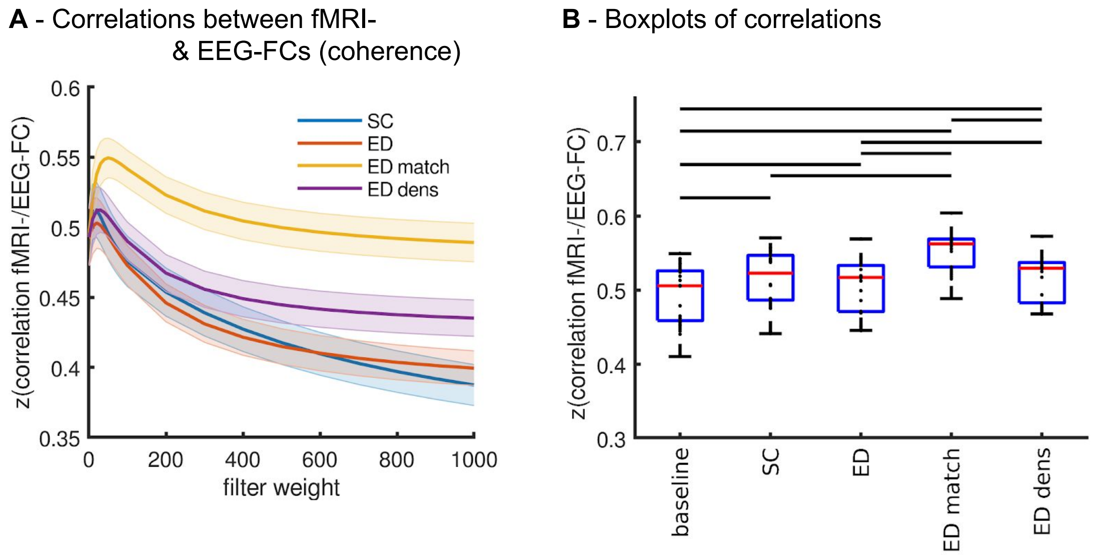


Figure S8: (related to Discussion) **A**:Fit (z-transformed correlation) between the EEG-FC (beta band, *coherence*) computed from time courses with different filter weights (G) and the fMRI-FC. The shaded regions mark the 95% confidence interval. **B**: Boxplots showing results of the Wilcoxon signed rank test comparing the maximum fits (shown in panel A) across versions of the SC as well as to the baseline correlation between unfiltered EEG-FC and fMRI-FC. Black bars mark significant differences. Red lines mark the median, each black dot marks the value for one subject.

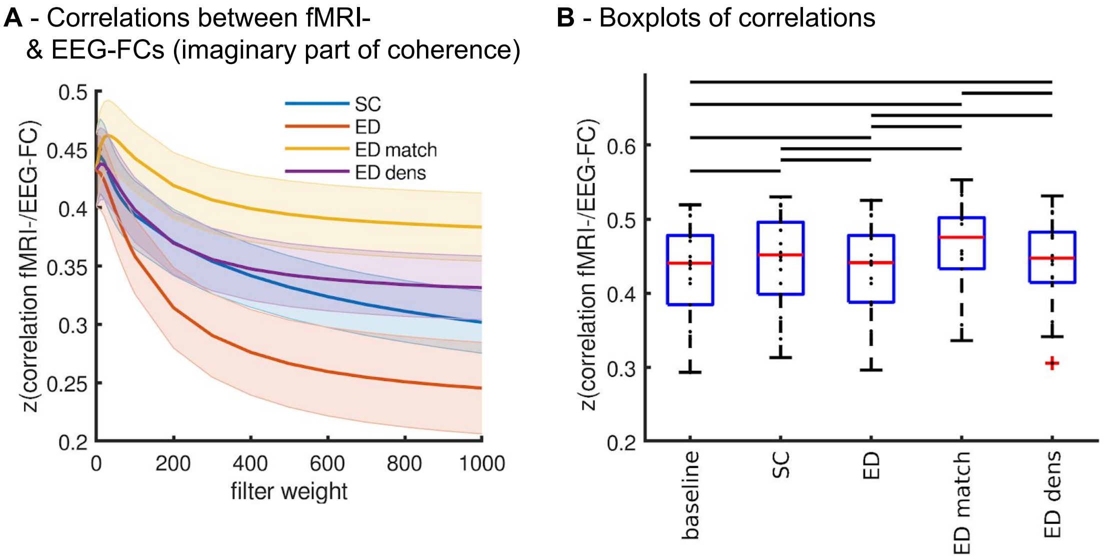


Figure S9: (related to Discussion) **A**:Fit (z-transformed correlation) between the EEG-FC (beta band, *imaginary part of coherence*) computed from time courses with different filter weights (G) and the fMRI-FC. The shaded regions mark the 95% confidence interval. **B**: Boxplots showing results of the Wilcoxon signed rank test comparing the maximum fits (shown in panel A) across versions of the SC as well as to the baseline correlation between unfiltered EEG-FC and fMRI-FC. Black bars mark significant differences. Red lines mark the median, each black dot marks the value for one subject.

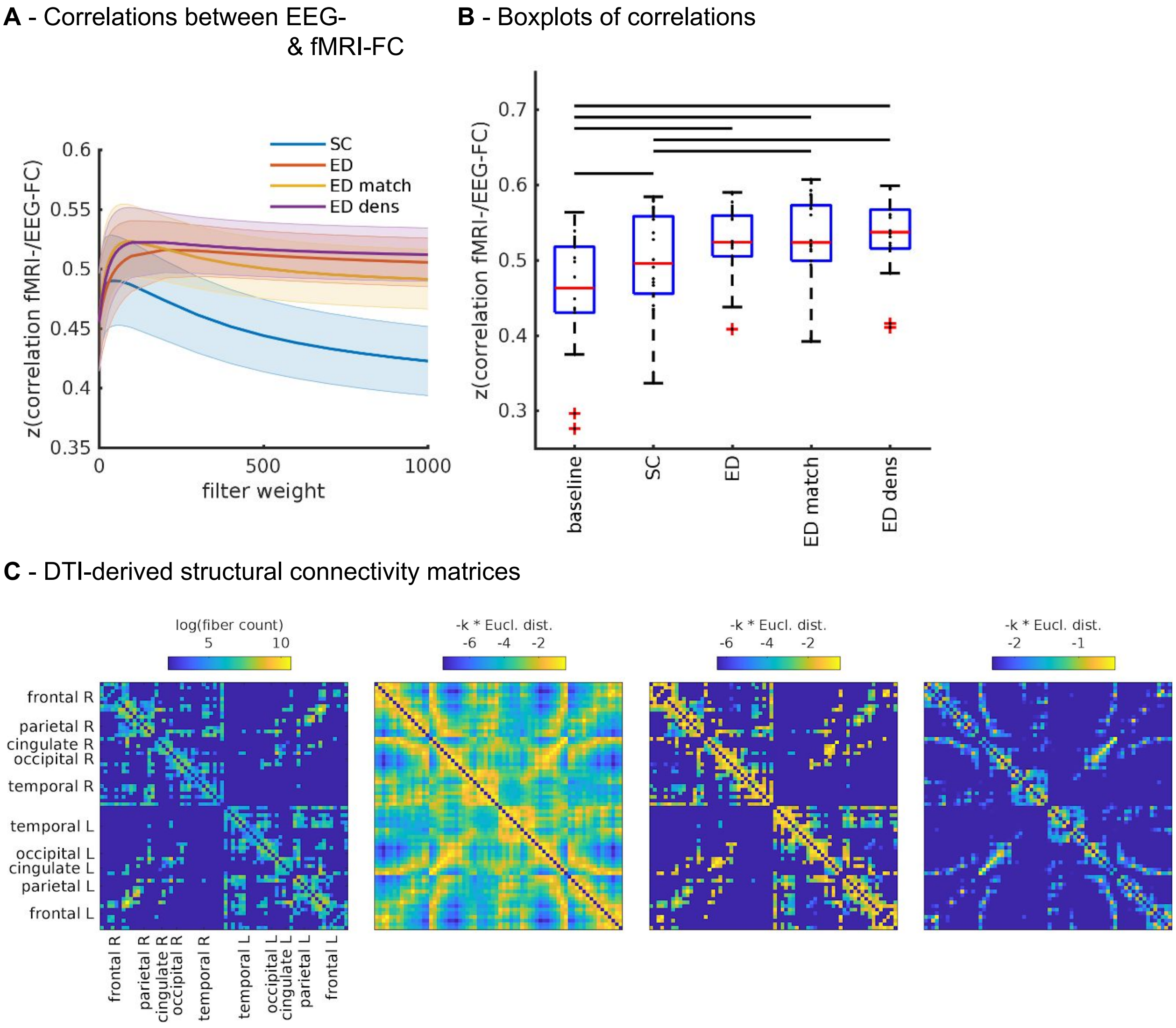


Figure S10: (related to Results - “Graph filtering increases resemblance between EEG-FC and fMRI-FC”, Figure 4) A: Fit (z-transformed correlation) between EEG-FCs (beta band) obtained using the DTI-derived SC-matrix and the fMRI-FC. The shaded regions mark the 95% confidence interval. **B**: Boxplots summarizing results of the Wilcoxon signed-rank test comparing individuals’ maximum fits (shown in panel A) across versions of the SC as well as to the baseline correlation between unfiltered EEG-FCs and fMRI-FC. Black bars mark significant differences. Red lines mark the median, each black dot marks the value for one subject. Note that we did not compare the medians, but the individual differences (see Methods).

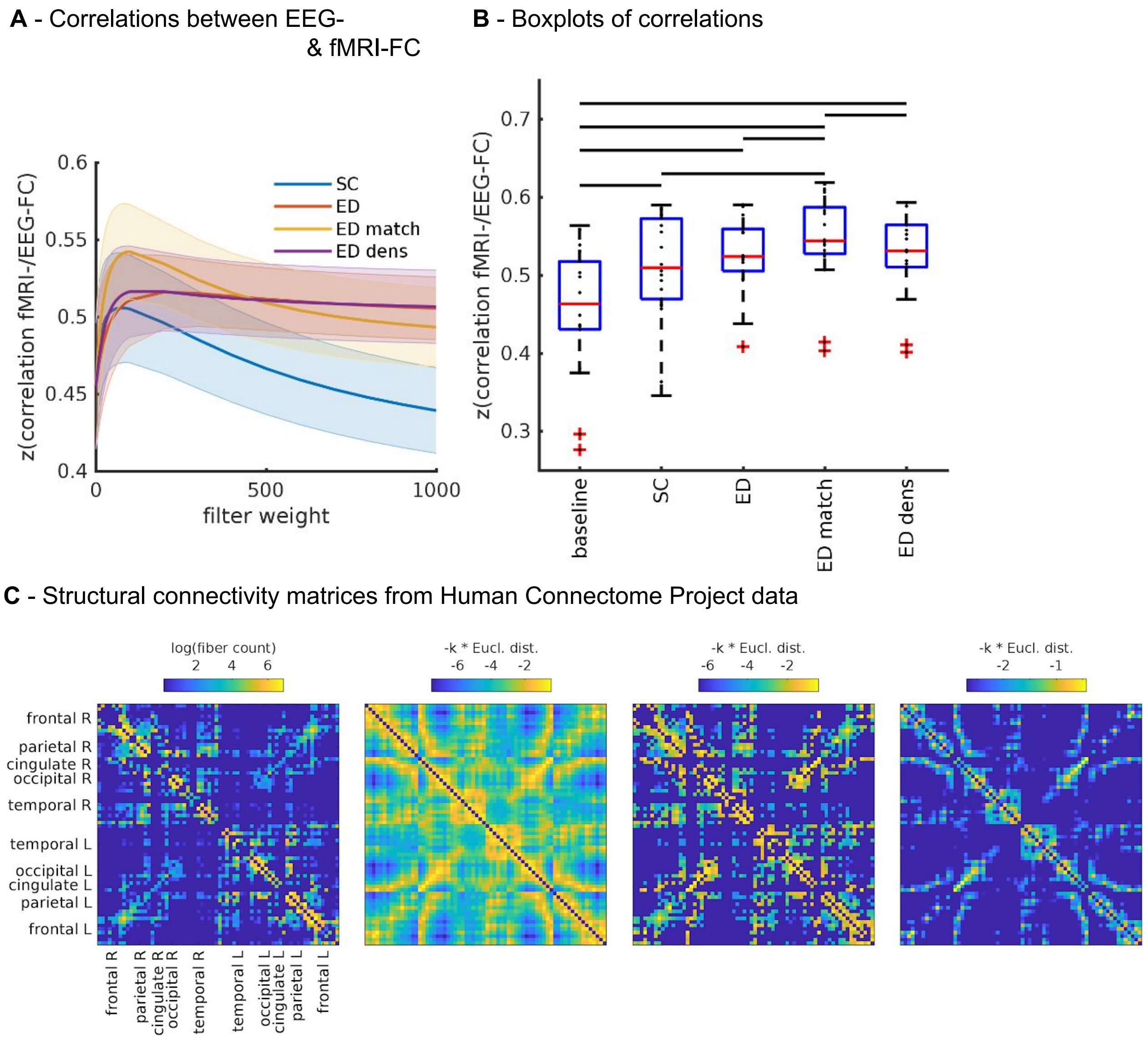


Figure S11: (related to Results - “Graph filtering increases resemblance between EEG-FC and fMRI-FC”, Figure 4) **A**: Fit (z-transformed correlation) between EEG-FCs (beta band) obtained using the **HCP-SC-matrix** and the fMRI-FC. The shaded regions mark the 95% confidence interval. **B**: Boxplots summarizing results of the Wilcoxon signed-rank test comparing individuals’ maximum fits (shown in panel A) across versions of the SC as well as to the baseline correlation between unfiltered EEG-FCs and fMRI-FC. Black bars mark significant differences. Red lines mark the median, each black dot marks the value for one subject. Note that we did not compare the medians, but the individual differences (see Methods).

**A** - Comparison between structural connectivity matrices

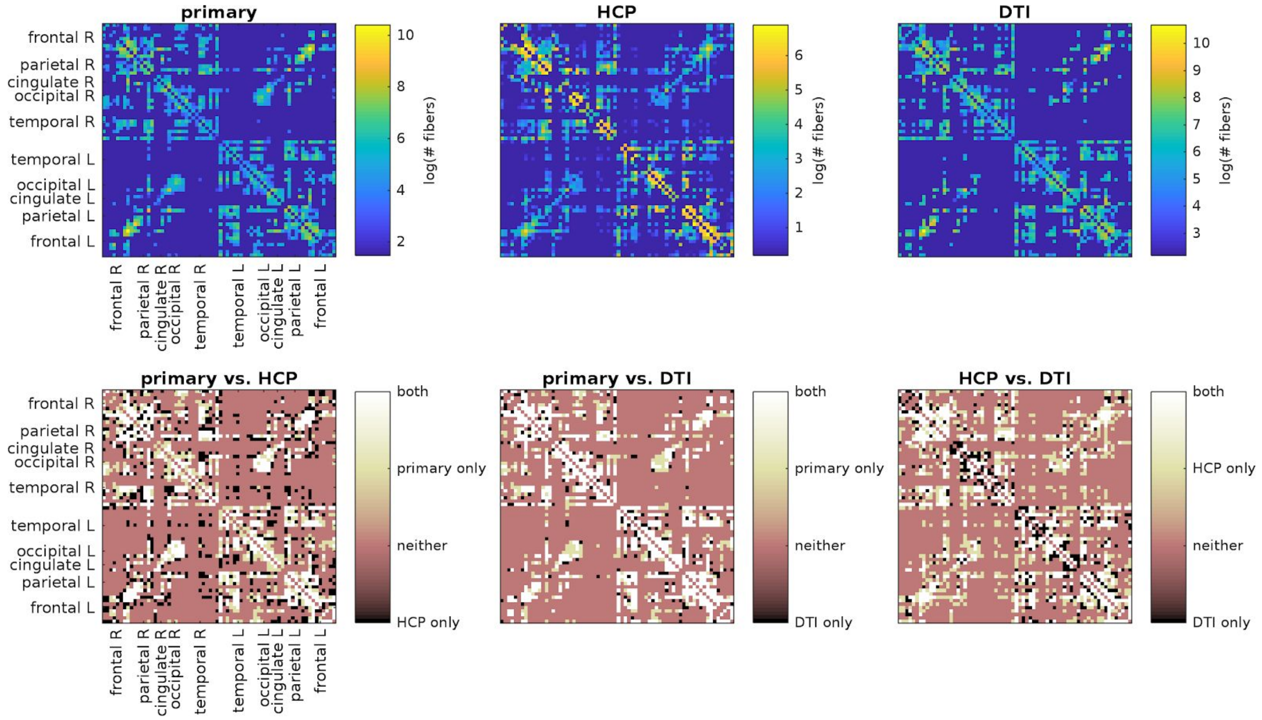


**B** - Effects of differences in structural connectivity matrices on the filtered EEG-FCs

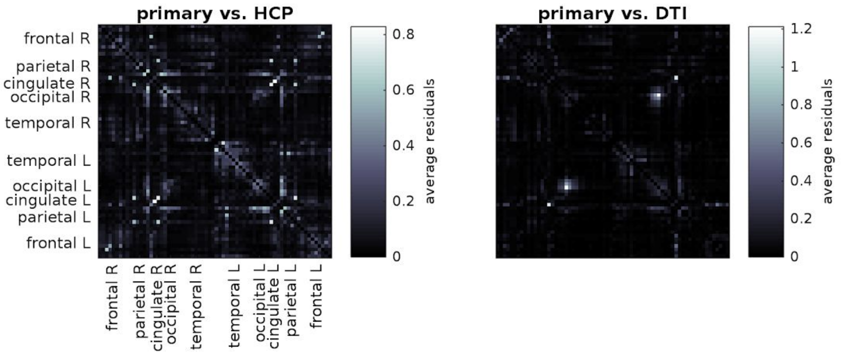


Figure S12: (related to Results - “Graph filtering increases resemblance between EEG-FC and fMRI-FC”) We repeated our main analysis (“primary”) with two structural connectivity matrices from two separate cohorts, “HCP”, a high-quality dataset from the Human Connectome Project, and “DTI”, a comparatively lower quality dataset acquired locally (see Methods for details). **A**: Each SC matrix (logarithm of the number of streamlines/fibers) is depicted in the top row, as well as comparisons between existing and absent fibers in the bottom row. While a core set of fibers is present in all SCs, there are also important differences. **B**: To test the effect of these differences on the resulting filtered EEG-FC matrices, we show the deviations (residuals) of each FC-pair from a perfect linear fit with the “primary” best EEG-FC (shown in Figure 5C). The largest deviations are located on the secondary diagonal, pointing to a lack in homotopic connections.

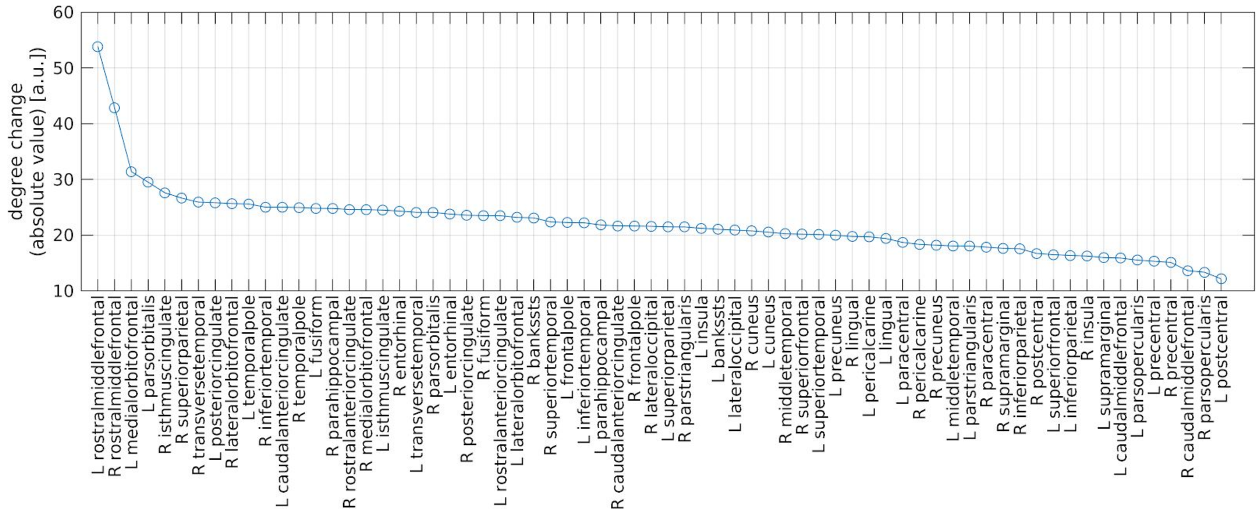


Figure S13: (related to Figure 6) Degree changes of all 68 ROIs induced by filtering. Average EEG-FCs of the beta band before and after filtering (G=100, “ED match”, best fit to fMRI-FC) were resampled and the absolute values of the difference was taken. The values plotted here are the sums over the rows/columns.

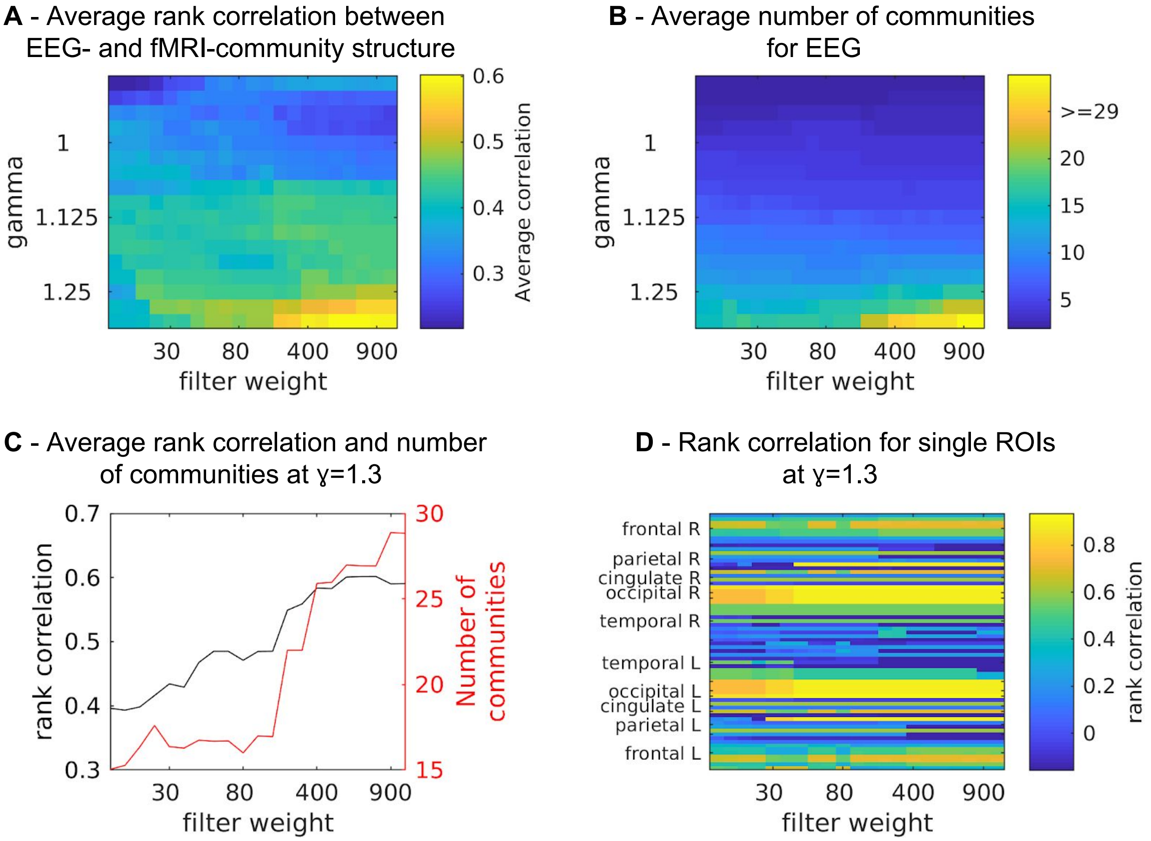


Figure S14: (related to Figure 9) Results of the community analysis when the number of communities is not limited to be no greater than 7. **A**: Agreement between community structure of EEG- and fMRI-FC as measured by average rank correlations between “community matrices” (not shown). In this case, the overall maximum of 0.60 in community agreement as measured by rank correlation between community matrices is found at G=700 and ɣ=1.3. **B**: Average number of communities found by the Louvain clustering algorithm. At G=700 and ɣ=1.3, the number of communities is 27. **C:** Rank correlations and number of communities for ɣ=1.3. **D**: Rank correlations between rows/columns of “community matrices” of EEG- and fMRI-FC, for ɣ=1.3.

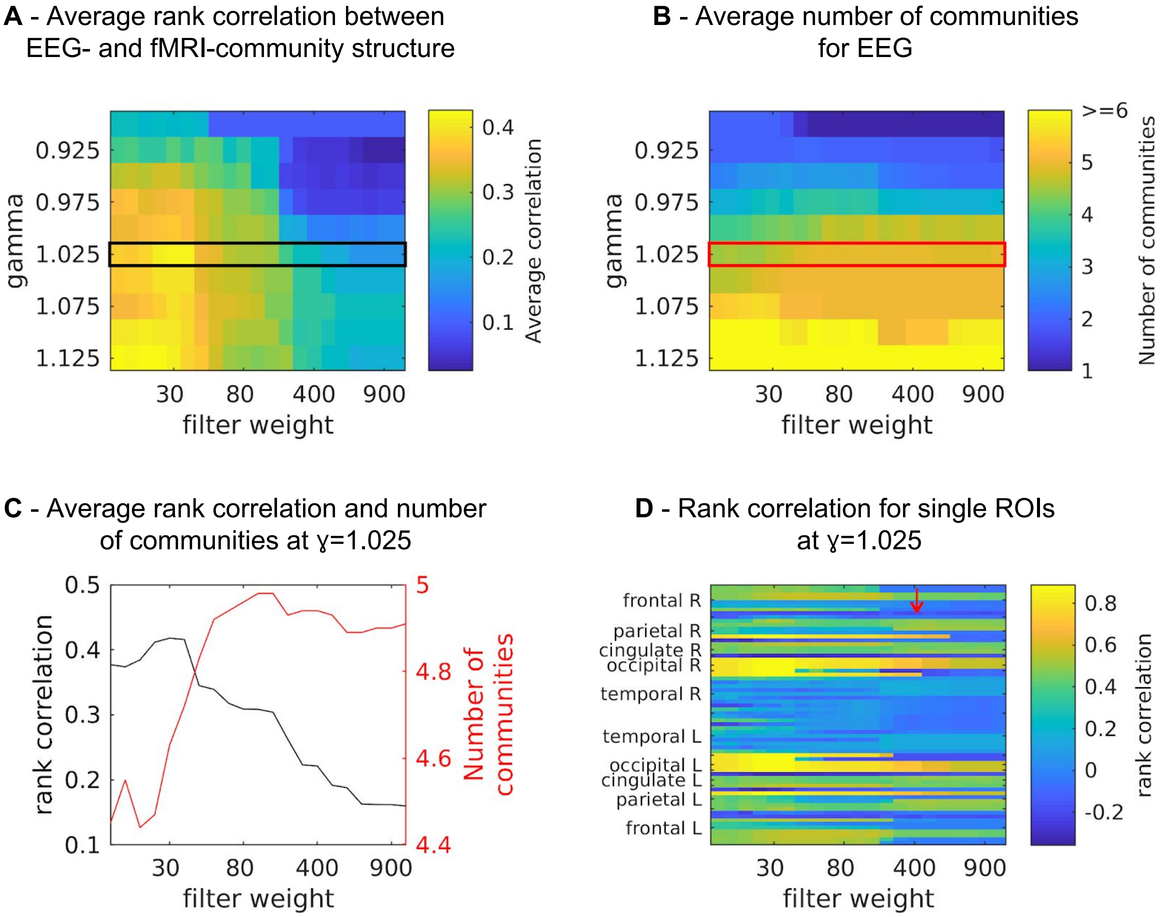


Figure S15: (related to Figure 9) Results of the community analysis when using the SC graph as a filter. **A**: Agreement between community structure of EEG- and fMRI-FC as measured by average rank correlations between "community matrices" (not shown). The black box marks the ɣ which is shown in panel C. **B**: Average number of communities found by the Louvain clustering algorithm. The red box marks the ɣ which is shown in panel C. **C:** Rank correlations and number of communities for ɣ=1.025 (marked in the same colors in panels A and B). **D**: Rank correlations between rows/columns of "community matrices" of EEG- and fMRI-FC, for ɣ=1.025 (marked with colored boxes in panels A and B). The red arrow marks the area for which rank correlations of parietal areas are increased - see Figure S5.

**A** - Community co-assignments of right paracentral lobule before filtering

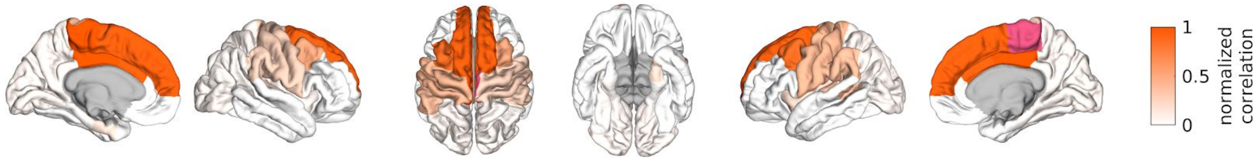


**B** - Community co-assignments of right paracentral lobule after filtering with the SC graph, G=400

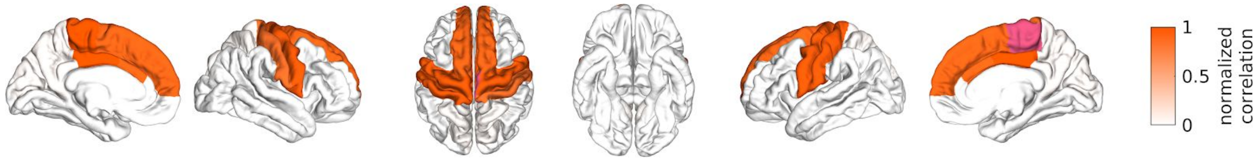


**C** - Community co-assignments of right paracentral lobule in fMRI

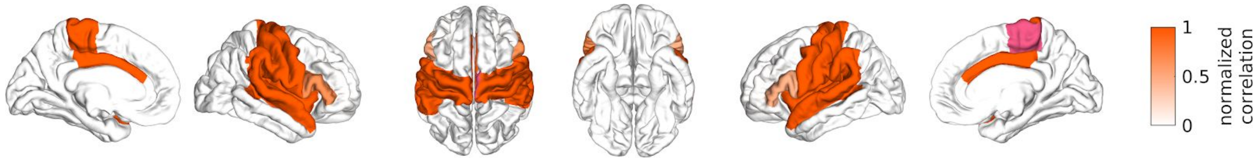


Figure S16: (related to Discussion) Community co-assignments of right paracentral lobule. **A**: Before filtering, this region is mostly grouped with some frontal regions. This figure looks different from what is shown for the same region in Figure 11A because the values of ɣ are different. **B**: After filtering *with the SC graph*, the region is robustly grouped with bilateral pre- and postcentral gyri, however, the grouping with the frontal regions is not removed. Most importantly, the overall fit to the fMRI community structure for these parameter settings (ɣ=1.025, G=400) is only 0.22. **C**: In fMRI, this network includes the insula as well as some temporal regions and is symmetrical.

Table S2: (related to Results: “Graph filtering increases resemblance between EEG-FC and fMRI-FC“, and Discussion: “Comparison to other methods that attenuate volume conduction”) Subject-wise correlations between EEG-FC and fMRI-FC with and without filtering as well as with orthogonalization (without filtering).

| S | 1 | 2 | 3 | 4 | 5 | 7 | 8 | 9 | 12 | 13 | 14 | 15 | 16 | 17 | 18 | 19 | 20 | 21 |
|---|---|---|---|---|---|---|---|---|---|---|---|---|---|---|---|---|---|---|
| unfiltered | 0.36 | 0.41 | 0.41 | 0.46 | 0.44 | 0.48 | 0.46 | 0.49 | 0.42 | 0.42 | 0.51 | 0.49 | 0.51 | 0.44 | 0.29 | 0.27 | 0.41 | 0.36 |
| filtered (best) | 0.50 | 0.52 | 0.51 | 0.54 | 0.50 | 0.56 | 0.54 | 0.54 | 0.48 | 0.51 | 0.55 | 0.56 | 0.53 | 0.54 | 0.41 | 0.40 | 0.50 | 0.51 |
| orthogonalized | 0.05 | 0.08 | 0.31 | 0.27 | 0.16 | 0.19 | 0.20 | 0.22 | 0.30 | 0.23 | 0.12 | 0.32 | 0.31 | 0.19 | 0.01 | 0.14 | 0.17 | 0.26 |

Table S3: (related to Methods-Data section) Number and lengths of artifact-free intervals that were included in the envelope correlation-based FC analysis, as well as the number of 3s-segments used for computing coherence and imaginary part of coherence, for each subject.

| subject ID | # intervals | lengths intervals (seconds, Fs=1000Hz) | | | | | | | | | | | | | # segments ([i]coh) |
|---|---|---|---|---|---|---|---|---|---|---|---|---|---|---|---|
| s01 | 9 | 52 | 36 | 42 | 37 | 27 | 26 | 26 | 29 | 20 | | | | | 64 |
| s02 | 6 | 90 | 20 | 22 | 20 | 31 | 39 | | | | | | | | 64 |
| s03 | 7 | 260 | 63 | 37 | 27 | 20 | 164 | 21 | | | | | | | 167 |
| s04 | 9 | 37 | 51 | 45 | 57 | 20 | 24 | 73 | 213 | 153 | | | | | 111 |
| s05 | 2 | 383 | 35 | | | | | | | | | | | | 97 |
| s07 | 5 | 96 | 38 | 80 | 39 | 29 | | | | | | | | | 63 |
| s08 | 5 | 60 | 39 | 92 | 329 | 48 | | | | | | | | | 155 |
| s09 | 7 | 33 | 47 | 46 | 54 | 38 | 38 | 70 | | | | | | | 77 |
| s12 | 6 | 77 | 72 | 92 | 79 | 162 | 253 | | | | | | | | 227 |
| s13 | 9 | 41 | 89 | 20 | 26 | 25 | 21 | 26 | 24 | 23 | | | | | 61 |
| s14 | 9 | 47 | 30 | 26 | 93 | 60 | 59 | 87 | 39 | 42 | | | | | 61 |
| s15 | 9 | 26 | 37 | 32 | 22 | 33 | 179 | 38 | 114 | 26 | | | | | 85 |
| s16 | 13 | 38 | 24 | 25 | 21 | 44 | 28 | 24 | 40 | 41 | 31 | 45 | 53 | 26 | 120 |
| s17 | 12 | 26 | 109 | 62 | 38 | 69 | 40 | 40 | 33 | 85 | 31 | 35 | 81 | | 160 |
| s18 | 2 | 25 | 20 | | | | | | | | | | | | 5 |
| s19 | 4 | 54 | 254 | 22 | 83 | | | | | | | | | | 96 |
| s20 | 7 | 68 | 24 | 23 | 27 | 39 | 44 | 19 | | | | | | | 19 |
| s21 | 8 | 64 | 93 | 117 | 48 | 23 | 19 | 26 | 40 | | | | | | 104 |
